# Pulmonary Fibrosis Enhances Vasodilation to Calcitonin Gene-Related Peptide

**DOI:** 10.64898/2026.05.10.724169

**Authors:** Charles E. Norton

## Abstract

**Background:** Calcitonin gene related peptide (CGRP) hyperpolarizes pulmonary arterial smooth muscle cells (SMCs) and endothelial cells (ECs) through PKA-dependent activation of K_ATP_ channels. CGRP can diminish the severity of pulmonary fibrosis (PF), however, the effects on vascular signaling were poorly defined. We hypothesized that hyperpolarization to CGRP would be augmented in a mouse model of PF.

**Methods:** PF was induced in male and female C57BL/6 mice by intratracheal delivery of bleomycin (3 wk), with saline used as control (sham). Pulmonary arteries (PAs; 100-150 µm diameter) were cannulated and pressurized to 16 cmH_2_O, and endothelial tubes were studied in complementary experiments to eliminate the influence of SMCs. Membrane potential (V_m_) was recorded continuously using intracellular microelectrodes. Responses were also evaluated in isolated lungs preconstricted with U46619 (∼10 mmHg).

**Results:** PF led to greater indices of PH in males vs. females. Isolated lungs and PAs from male PF mice had enhanced vasodilation and hyperpolarization of V_m_ to CGRP, although no effect was observed in females. The greater vasodilation and hyperpolarization of SMCs to CGRP in males persisted in endothelium-disrupted PAs and during treatment with L-NAME indicating that ECs are not required for greater responsiveness to CGRP. With no effect on resting V_m_, inhibition of K_ATP_ channels or PKA significantly attenuated hyperpolarization of SMCs and ECs, attenuated vasodilation to CGRP in PAs, and eliminated differences between groups in males. Direct activation of PKA, but not K_ATP_, evoked greater V_m_ hyperpolarization and vasodilation in PF vs. sham PAs and lungs. Although no difference in sensory nerves was observed in fibrotic mice, perivascular nerve stimulation evoked greater vasodilation in PAs.

**Conclusions:** In a mouse model of PF, CGRP-dependent hyperpolarization of pulmonary arterial SMCs and ECs is augmented through increased PKA-dependent activation of K_ATP_ channels leading to increased vasodilator sensitivity.

## INTRODUCTION

Pulmonary fibrosis (PF) is a debilitating disease associated with dyspnea, cough, and impaired gas exchange resulting from excessive deposition of extracellular matrix component^1^. It can result from occupational exposure (e.g., asbestos), acute lung injury, or idiopathic causes. The incidence of PF has risen in recent decades likely relating to increased exposures to smoke and particulate matter associated with modern urban lifestyles^2^. The average life expectancy for an individual after being diagnosed with PF is 3 to 5 years^3^. While there is no cure for the disease, therapeutic treatments can slow its progression. The presence of pulmonary hypertension (PH) in PF significantly increases mortality^4^ and develops over time in most patients with PF^4, 5^. Although most cases of PH in PF are mild or moderate, many patients can have severe PH^5^. Systolic pulmonary arterial pressure (PAP) has a strong inverse correlation with survival. Specifically, patients with PAP > 50 mmHg have a survival of < 1 year compared to a survival of > 4 years in patients with PAP below 50 mmHg^6^.

Despite the key role of PH in PF, little direct attention has been paid to the processes which elicit vascular dysfunction. Historically, mechanisms of PF-induced PH have been thought to mimic chronic hypoxia, however, emerging evidence suggests that additional signaling events are involved^6, 7^. A particular gap in knowledge exists with regards to regulation of pulmonary vascular tone in the context of interstitial lung diseases such as PF. Administration of vasodilators is an integral tool for the treatment of PH^8^, however, as the sensitivity to a variety of vasodilators is impaired in PH^9^, identifying those which remain effective under conditions of vascular dysfunction in PF is important to improve therapeutic interventions.

Sensory nerves are located throughout the pulmonary vascular tree^10^ and regulate the diameter of resistance arteries. Calcitonin gene-related peptide (CGRP) is the primary neurotransmitter released from sensory nerves and is a potent vasodilator and anti-inflammatory agent^11-13^. CGRP activates G protein-coupled receptors on both smooth muscle cells (SMCs) and endothelial cells (ECs) to elicit vasomotor effects^11, 14^. In chronic hypoxia-induced PH, there is reduced vasodilator sensitivity to CGRP^15^, however, CGRP signaling can diminish the severity of PF^16, 17^. Therefore, we hypothesized that CGRP-dependent dilation would be augmented in pulmonary arteries (PAs) during the initial stages of PF. The present findings reveal that three weeks following the initiation of PF with bleomycin in mice, PH is more severe in males vs. females, and PAs from males have greater vasodilation to sensory nerve stimulation mediated by CGRP.

## METHODS

All protocols and experimental procedures were reviewed and approved by the University of Missouri Animal Care and Use Committee (Protocol #40469). Detailed Materials and Methods are described in the Supplemental Material and Major Resource Table. Data supporting the findings of this study are available at https://doi.org/10.7910/DVN/GT07XB.

## RESULTS

### Bleomycin Induces Pulmonary Fibrosis and Pulmonary Hypertension

Intratracheal delivery of bleomycin elicited a notable increase in collagen deposition in lungs from males and females after 3 wk (Figures 1A and B; S1) indicating the development of PF. This corresponded to an increase in cellularity in lung sections from both sexes (Figure 1C). Mice exposed to bleomycin had increased RV systolic pressure (RVSP; Figure 1D and E) with no differences in anesthetized heart rate (male sham = 301±24, male bleomycin = 296±18, female sham = 304±21, female bleomycin = 291±15, beats per minute) indicating the development of PH. In addition, PF resulted in right ventricular (RV) hypertrophy as indicated by an increase in Fulton index (Figure 1F). However, both indices of PH were significantly reduced in females as compared to males similar to what is observed in response to hypoxia-induced PH^18^. RV/body weight ratios were similarly elevated without a significant effect on left ventricular weight (Table S1). Male mice with PF-induced PH exhibited lower body weight when compared to their sham controls (P=0.11 for females). PF led to an increase in hematocrit which was greater in males vs. females (Table S1).

**Figure 1.**
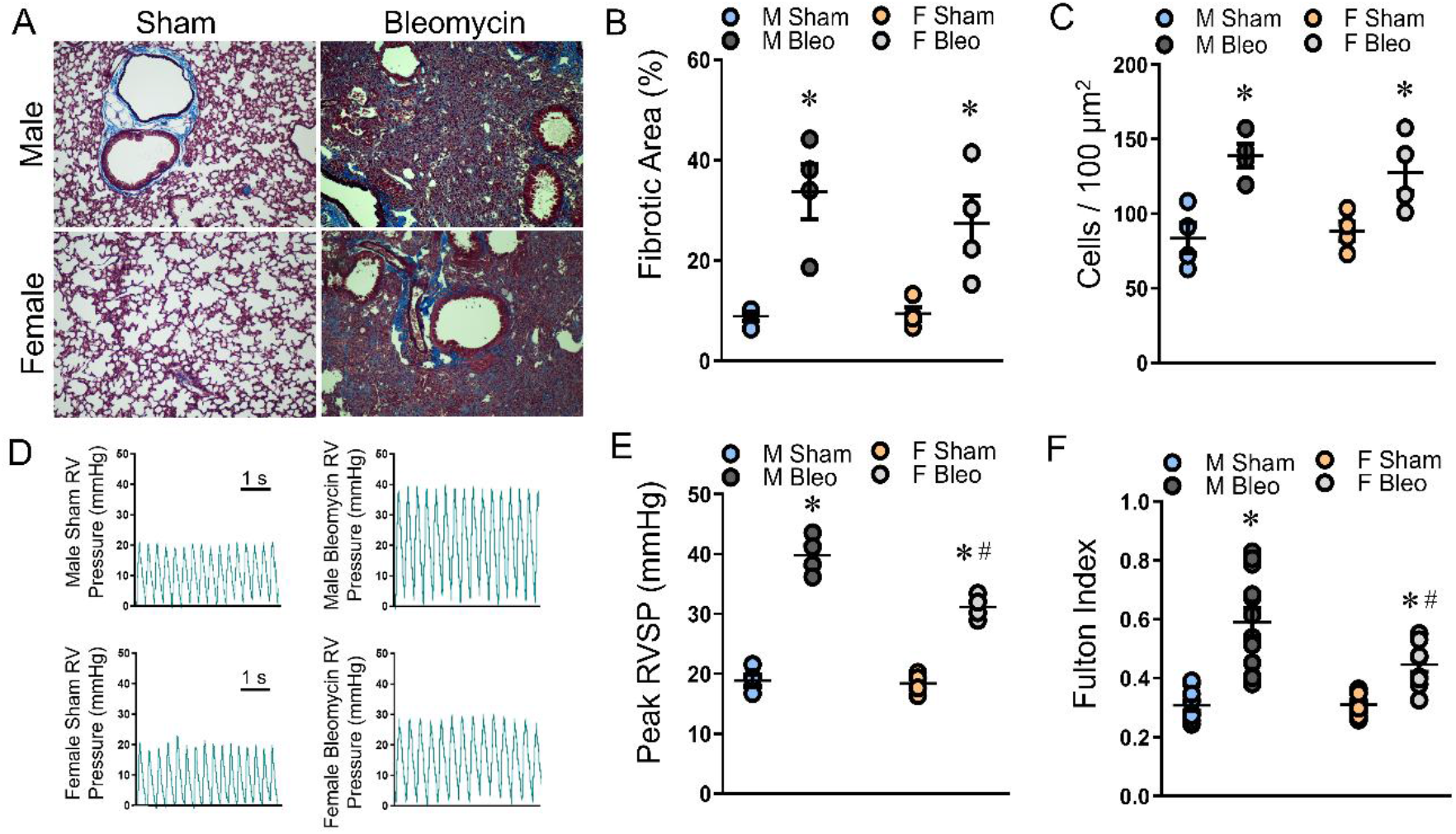
Bleomycin induces PF and Pulmonary Hypertension. A) Representative Masson’s Trichrome staining in Sham and Bleomycin treated lungs from male (M) and female (F) mice at 10× magnification (scale bar = 200 µm). B) Quantification of fibrotic area (% collagen) in lung sections from males and females. C) Cellularity (counted nuclei) in lungs from males and females. D) Sample traces of right ventricular (RV) pressure in anesthetized sham and bleomycin treated mice. E) Summary data for peak RV systolic pressure (RVSP) and F) right ventricular hypertrophy (Fulton’s index) as indices of pulmonary hypertension following bleomycin treatment in males and females. Values are means ± SEM; n=4-10/group. *P<0.05 bleomycin vs. sham. ^#^P<0.05 female vs. male.

### Pulmonary Fibrosis Enhances Vasodilation to CGRP

Baseline and maximal diameters and vessel wall [Ca^2+^]_i_ were similar between isolated PAs pressurized to 16 cmH_2_O (∼12 mmHg) from sham and bleomycin-treated mice in males and females (Table S2). Due to the minimal basal (spontaneous) tone in PAs studied at 16 cmH_2_O, vessels were preconstricted with uridine triphosphate (UTP). Vasoconstriction to UTP and the corresponding [Ca^2+^]_i_ response was unaltered by bleomycin treatment in PAs from either males or females (Figure S2). In vessels preconstricted with an EC_50_ value of UTP (5×10^-6^ M), CGRP (10^-10^-10^-6^ M) evoked a concentration-dependent dilation in each group. For male PAs, vasodilation to CGRP was augmented in mice treated with bleomycin (Figure 2A) with a greater reduction in vessel wall [Ca^2+^]_i_ (Figure 2B). In PAs from females, vasodilation to CGRP and the corresponding [Ca^2+^]_i_ responses were not significantly altered by PF.

**Figure 2.**
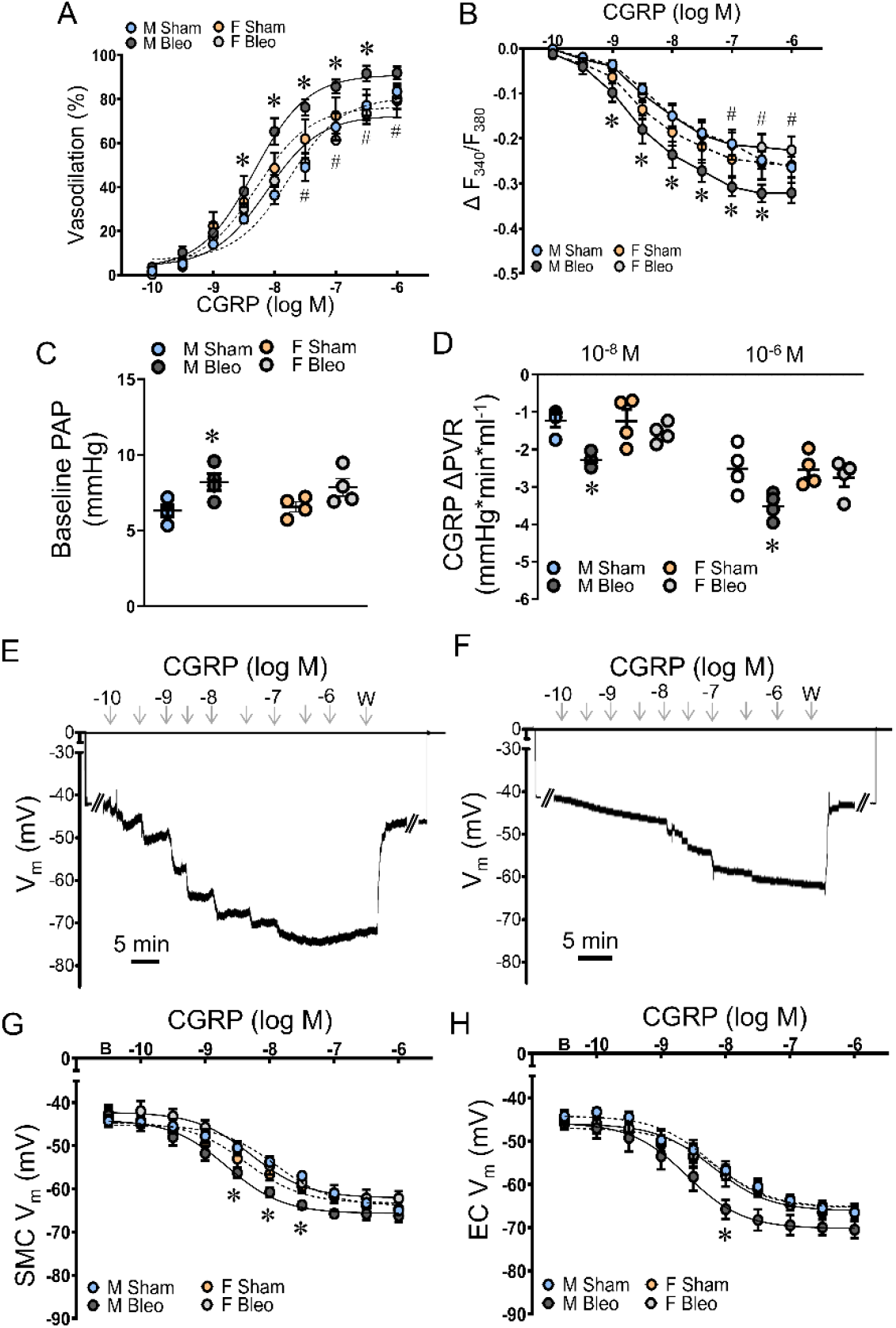
PF enhances pulmonary vasodilator sensitivity to CGRP in males. A) Vasodilation to CGRP (10^-10^-10^-6^ M) and B) corresponding Ca^2+^ responses (ΔF_340_/F_380_) in isolated PAs preconstricted with UTP from male and female mice. C) Pulmonary arterial pressures (PAP) at baseline and D) changes in pulmonary vascular resistance (PVR) in response to CGRP (10^-8^-10^-6^ M) in isolated lungs preconstricted with U46619. Representative continuous recordings of membrane potential (V_m_) in a PA from bleomycin male (E) and an endothelial tube from sham female (F) mice illustrate concentration-dependent hyperpolarization of SMCs and ECs. Note sharp change in V_m_ upon electrode entry and exit from each cell; W, washout. Summary data for V_m_ of (G) SMCs from intact PAs and (H) ECs from endothelial tubes to CGRP in male and female mice following bleomycin or sham treatment. B, baseline V_m_. Values are means ± SEM; n=4-5/group. *P<0.05 bleomycin vs. sham. ^#^P<0.05 female vs. male in bleomycin.

To evaluate changes in vascular function in the whole lung, we performed experiments in ventilated, isolated saline perfused lungs. Consistent with reduced airway compliance, peak tracheal pressures during ventilation were higher in bleomycin vs. sham lungs (Table S3). Baseline peak PAP was augmented in isolated lungs from male, but not female (P=0.10) mice treated with bleomycin (Figure 2C). In lungs preconstricted with U46619 (∼10 mmHg; Table S3), CGRP led to a greater reduction of pulmonary vascular resistance (PVR) in bleomycin-treated lungs from males but not females (Figure 2D).

### Pulmonary Fibrosis Enhances Hyperpolarization to CGRP

Initial resting potential was similar for across preparations and treatments (Table S4). During continuous recording of V_m_, CGRP (10^-10^-10^-6^ M) hyperpolarized SMCs and ECs from males and females (Figures 2E-D). For vessels from males, hyperpolarization was greater in bleomycin vs. sham mice. In females, hyperpolarization was similar between treatments and comparable in magnitude to sham males. In endothelial tubes isolated from male PAs, hyperpolarization to CGRP was augmented at a single concentration. There were no differences in EC hyperpolarization to CGRP in females. EC_50_ values for PF males were significantly lower than corresponding females or sham animals (Table S4).

### Enhanced Vasodilation to CGRP persists at Elevated Pressure

Given that greater vasodilation to CGRP was selectively observed in vessels from male mice exposed to bleomycin, we performed our subsequent experiments evaluating the mechanisms of this difference selectively in males. To determine whether the greater dilation to CGRP in males also occurred at higher pressures which occur during PH, experiments were repeated in PAs pressurized to 40 cmH_2_O (∼30 mmHg). Enhanced dilation to CGRP in PAs from PF mice remained at high pressure and there was a greater fall in vessel wall [Ca^2+^]_i_ in vessels from bleomycin vs. sham treated mice (Figure S3). For SMCs of isolated PAs, hyperpolarization to CGRP was enhanced at 40 cmH_2_O intraluminal pressure (Figure S4).

### Contribution of ECs to Enhanced Vasoconstriction to CGRP during Pulmonary Fibrosis

We further sought to evaluate the contribution of the endothelium to CGRP-dependent hyperpolarization and vasodilation in PAs from males. In endothelium disrupted PAs, or vessels treated with the NO signaling inhibitor LNAME (10^-4^ M), enhanced SMC hyperpolarization to CGRP persisted following bleomycin treatment (Figure S5). Furthermore, vasodilation, and [Ca^2+^]_i_ responses to CGRP remained augmented in mice exposed to bleomycin as compared to sham controls (Figure S6). Endothelial [Ca^2+^]_i_ responses to CGRP in endothelial tubes were unaltered by pulmonary fibrosis (Figure S7). Collectively these findings indicate that ECs are not required for the greater vascular SMC sensitivity to CGRP.

### K_ATP_ Channels Mediate Greater Hyperpolarization and Vasodilation to CGRP During Pulmonary Fibrosis

K_ATP_ channels are the primary mediators of vasodilation to CGRP in PAs^14^, however in systemic vessels, K_Ca_ channels can contribute to this response^19, 20^. Therefore, we sought to determine the respective contribution of potassium channels to mediate enhanced vasodilation to CGRP in vessels from males during PF. Within isolated lungs, glibenclamide nearly abolished the reduction of PVR during exposure to CGRP and eliminated differences between sham and bleomycin lungs (Figure 3A). Inhibition of K_ATP_ channels with glibenclamide (10^-6^ M)^14^ greatly reduced vasodilation to CGRP in preconstricted PAs (Figure 3B) and similarly limited the reduction in vessel wall Ca^2+^ (Figure 3C) for both bleomycin and sham treated mice nearly eliminating both responses. Consistently, K_ATP_ ^-/-^ mice (K_IR_ 6.1 global knockout) had greatly attenuated responsiveness to CGRP in isolated PAs from both groups (Figure 3D and E). Both pharmacological and genetic disruption of K_ATP_ signaling eliminated differences between bleomycin and sham mice, suggesting that greater vasodilation during PF is mediated by K_ATP_ channels. Furthermore, bleomycin-induced increases in RVSP and Fulton index were significantly enhanced in K_ATP_ ^-/-^ mice (Figure 3F and G) suggesting that K_ATP_ channel function limits the severity of PH resulting from PF.

**Figure 3.**
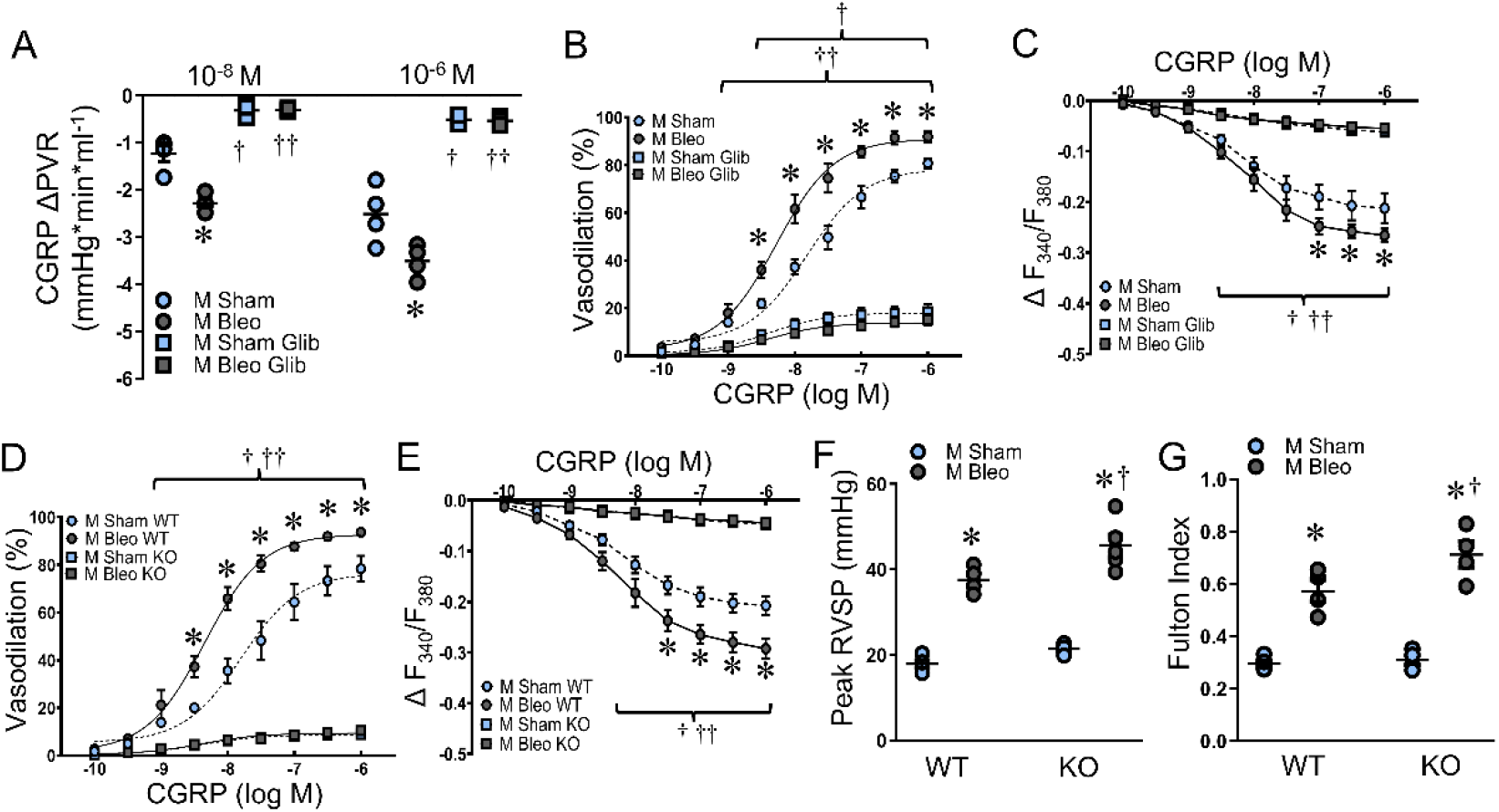
Loss of K_ATP_ function eliminates enhanced vasodilation between PF and sham mice. A) Changes in PVR to CGRP (10^-8^-10^-6^ M) in isolated lungs preconstricted with U46619 in the presence or absence of glibenclamide. B) Vasodilation and C) Ca^2+^ responses to CGRP in isolated pressurized PAs from males in the presence or absence of glibenclamide (10^-6^ M; Glib). D) Vasodilation and E) Ca^2+^ responses to CGRP in isolated pressurized PAs from K_ATP_ knockout (KO) mice or wildtype littermates (WT). F) RVSP and G) Fulton’s index in male mice from K_ATP_ knockout mice and wildtype littermates. Values are means ± SEM; n=4-5/group. *P<0.05 bleomycin vs. sham. ^†^P<0.05 glibenclamide/KO vs. vehicle/WT in sham. ^††^P<0.05 glibenclamide/KO vs. vehicle/WT in bleomycin.

Glibenclamide nearly abolished hyperpolarization to CGRP in both SMCs and ECs (Figures 4A-C) and eliminated the greater change in V_m_ in PAs from bleomycin mice. As the maximal V_m_ in these experiments was ≤ 5 mV, electrode placement in these experiments was confirmed with ACh (10^-5^ M) for ECs and KCl (10^-1^ M) for SMCs and these responses did not differ between groups (Table S5). Furthermore, hyperpolarization of SMCs and ECs was greatly attenuated in PAs from both sham and PF mice lacking K_ATP_ channels (Figures 4D-F).

**Figure 4.**
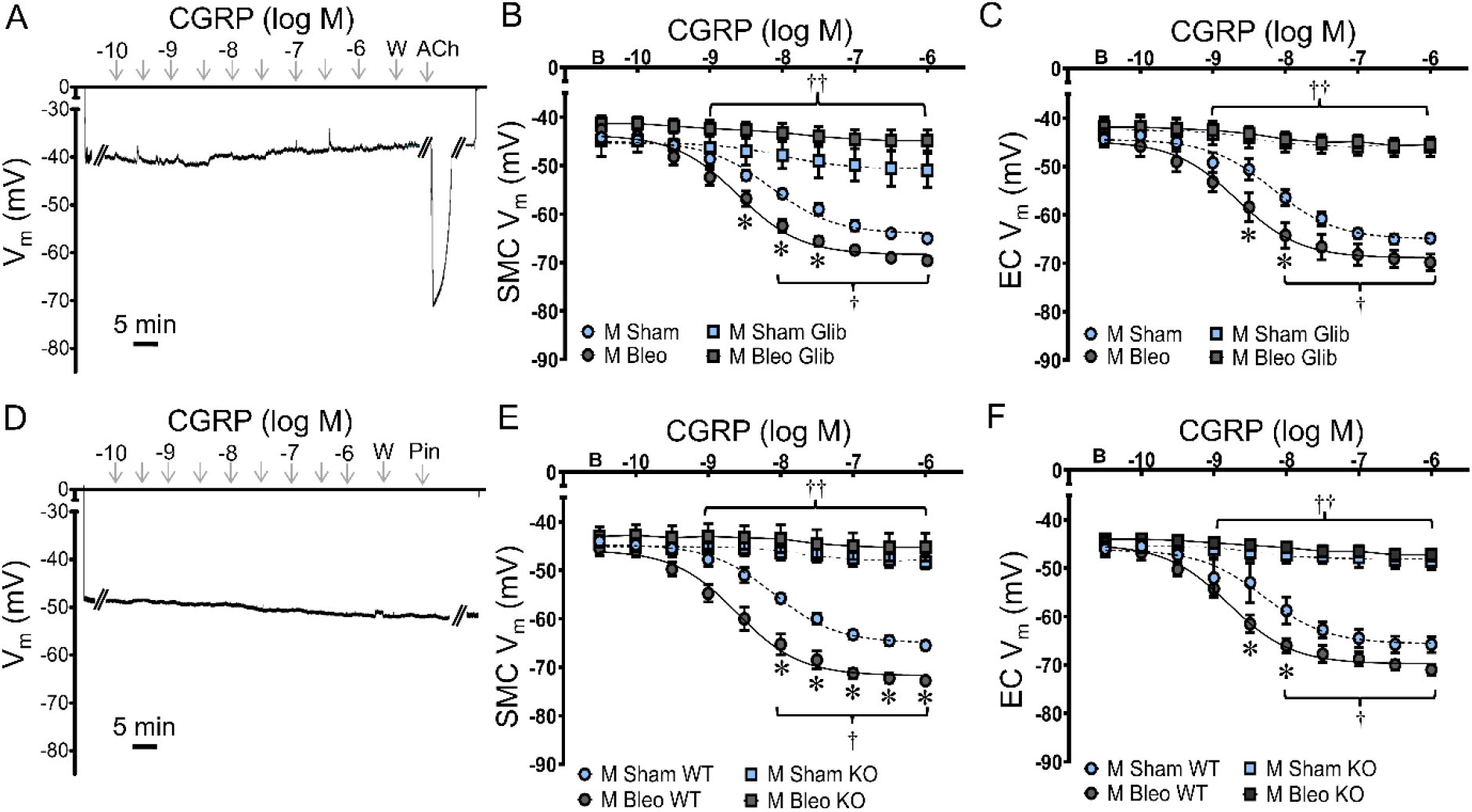
K_ATP_ channels contribute to greater hyperpolarization to CGRP in PF. A) Representative membrane potential (V_m_) recording in an EC tube from a male mouse treated with bleomycin. Summary data for V_m_ of B) SMCs and C) ECs in the presence or absence of glibenclamide (10^-6^ M; Glib). D) Representative membrane potential (V_m_) recording in an EC tube from a sham male mouse. Note lack of membrane potential response to the K_ATP_ agonist pinacidil (Pin; 10^-5^ M). Summary data for V_m_ of E) SMCs and F) ECs in the K_ATP_ knockout mice (KO) or wildtype littermates (WT). Values are means ± SEM; n=5-6/group. *P<0.05 bleomycin vs. sham. ^†^P<0.05 glibenclamide/KO vs. vehicle/WT in sham. ^††^P<0.05 glibenclamide/KO vs. vehicle/WT in bleomycin.

To further characterize the role of K_ATP_ channels in pulmonary vasodilation in PF, we directly activated K_ATP_ channels with pinacidil (10^-9^-10^-5^ M)^14^. Pinacidil evoked a concentration-dependent hyperpolarization in SMCs from intact arteries and ECs in endothelial tubes (Figure 5A-C). For both cell types, hyperpolarization to pinacidil did not differ between sham vs. bleomycin treatments with a similar potency (EC_50_: SMCs, bleomycin = 60 nM, sham = 66 nM; ECs, bleomycin = 106 nM, sham = 114 nM). In addition, pinacidil-dependent dilation was similar in isolated, pressurized PAs from sham and bleomycin mice (Figure 5D) with no differences in the [Ca^2+^]_i_ response between groups (Figure 5E). Consistently, pinacidil led to similar changes in PVR in sham and bleomycin treated isolated lungs (Figure 5F).

**Figure 5.**
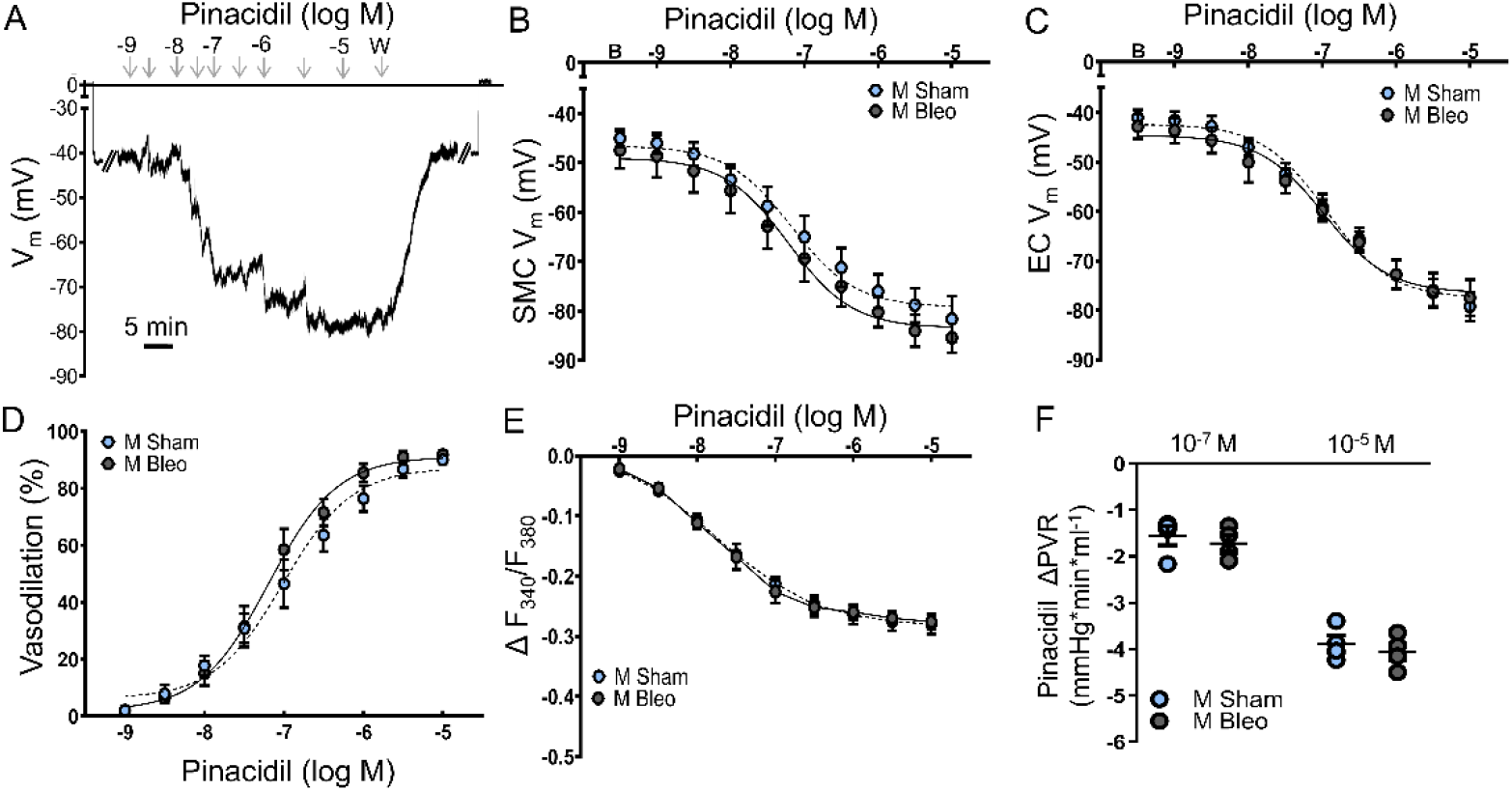
K_ATP_ channel activation does not evoke greater hyperpolarization or vasodilation. A) Representative membrane potential (V_m_) recording showing hyperpolarization to pinacidil (10^-9^-10^-5^ M) in an EC tube from a male mouse treated with bleomycin. Summary data for changes in V_m_ in response to pinacidil in B) SMCs and C) ECs. D) Vasodilation and E) Ca^2+^ responses to pinacidil in isolated pressurized PAs from sham and bleomycin treated males. F) Changes in PVR in response to pinacidil (10^-7^-10^-5^ M) in isolated lungs preconstricted with U46619. Values are means ± SEM; n=4-5/group. No significant differences were observed.

We also investigated the potential for K_Ca_ channels to mediate greater effects of CGRP within PAs with charybdotoxin (1×10^−7^ M; inhibits BK and IK) + apamin (3×10^−7^ M; inhibits SK)^21^. Combined inhibition of K_Ca_ channels did not significantly alter vasodilation or [Ca^2+^]_i_ responses to CGRP in PAs from either bleomycin or sham mice (Figure S8) and effects of PF to augment responsiveness to CGRP remained during K_Ca_ inhibition. K_Ca_ inhibition also did not alter CGRP-dependent hyperpolarization in SMCs and ECs in either bleomycin or sham treated mice (Figure S9).

### Role for PKA in Greater Hyperpolarization and Dilation to CGRP in PF

As K_ATP_ channels mediated, but did not directly evoke greater vasodilation in PAs from PF mice, we sought to evaluate differences in PKA-dependent vasodilation and hyperpolarization as it acts as an upstream target of K_ATP_ activation in PAs^14^. Inhibition of PKA with protein kinase inhibitor (14-22) amide (PKI, 10^-5^ M), used for its selectivity for PKA over PKG^22^, nearly eliminated vasodilation to CGRP in PAs preconstricted with UTP (Figure 6A), greatly attenuated the reduction in vessel wall [Ca^2+^]_i_ (Figure 6B), and eliminated differences between sham and bleomycin mice. In isolated lungs, PKI significantly reduced changes in PVR in response to CGRP and eliminated differences between sham and bleomycin mice (Figure 6C). In the presence of PKI, the maximal ΔV_m_ to CGRP was reduced to ≤5 mV in SMCs of intact arteries (Figure 6D and E), and differences were observed between sham and bleomycin groups. Similarly, PKA inhibition nearly abolished hyperpolarization to CGRP in endothelial tubes from sham and bleomycin mice (Figure 6F).

**Figure 6.**
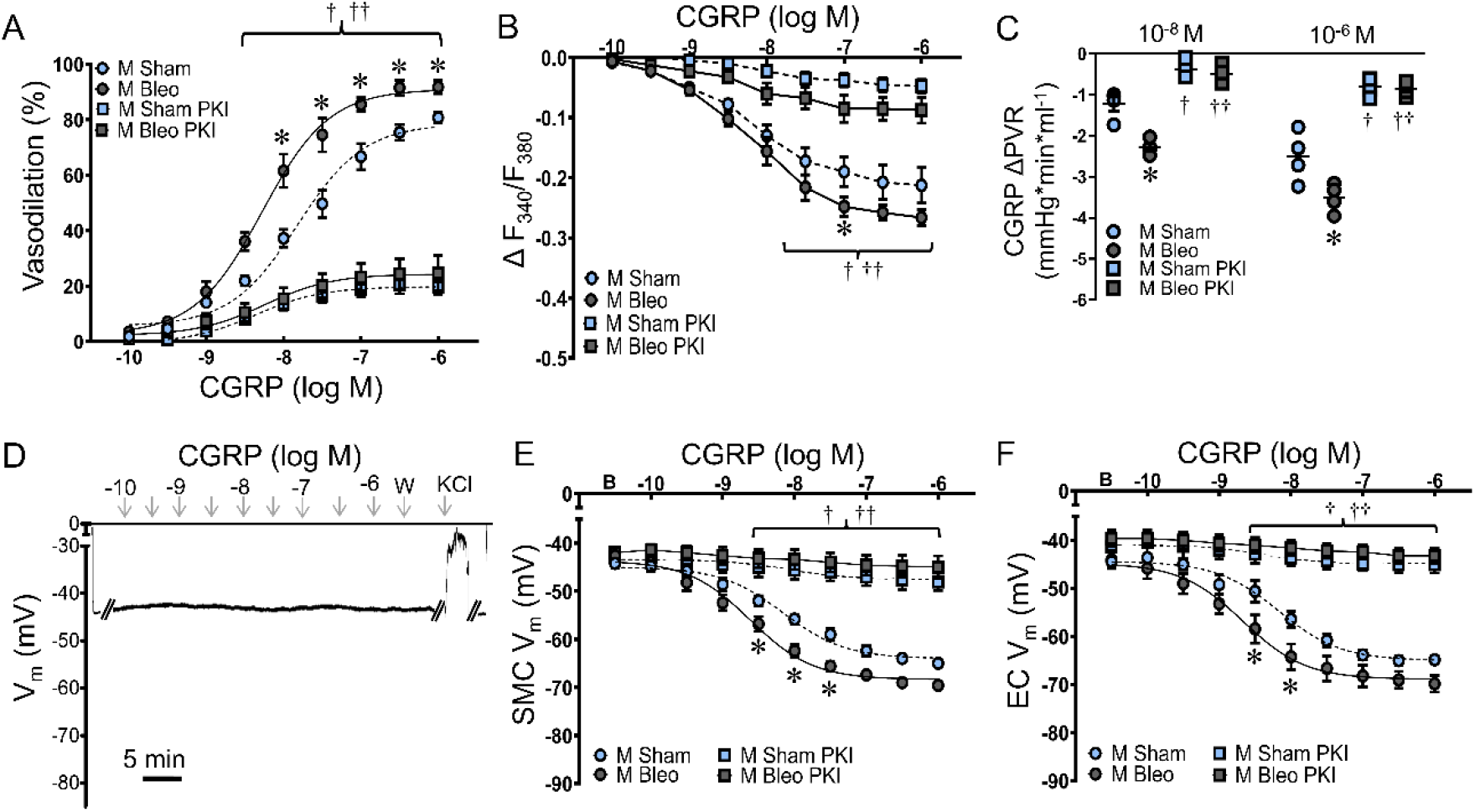
PKA mediates enhanced vasodilation and hyperpolarization to CGRP in PAs from PF mice. A) Vasodilation and B) Ca^2+^ responses to CGRP in isolated pressurized PAs from males in the presence or absence of PKI (10^-5^ M). C) Changes in PVR to CGRP (10^-8^-10^-6^ M) in isolated lungs preconstricted with U46619 in the presence or absence of PKI. D) Representative membrane potential (V_m_) recording in a SMC from a male PA from a bleomycin-treated mouse. Summary data for V_m_ of E) SMCs and F) ECs in the presence or absence of PKI. Values are means ± SEM; n=4-5/group. *P<0.05 bleomycin vs. sham. ^†^P<0.05 PKI vs. vehicle in sham. ^††^P<0.05 PKI vs. vehicle in bleomycin.

To determine whether PF alters PKA activation in PAs from males, we stimulated PKA with forskolin (10^-9^-10^-5^ M)^23^. Forskolin evoked a concentration-dependent vasodilation in preconstricted PAs from both groups of males, however, forskolin-dependent dilation was greater in bleomycin vs. sham PAs (Figure 7A). PKA activation simultaneously led to a more robust fall in [Ca^2+^]_i_ for PAs from PF mice (Figure 7B). In isolated lungs, forskolin led to a more robust fall in PVR in lungs from bleomycin mice compared to sham controls (Figure 7C). Consistently, forskolin evoked greater hyperpolarization of SMCs and ECs in bleomycin vs. sham PAs and endothelial tubes (Figure 7D-F). To verify effects of forskolin were mediated primarily through PKA signaling, we confirmed that >80% of forskolin-induced vasodilation and hyperpolarization was prevented in PAs treated with PKI (Figure S10).

**Figure 7.**
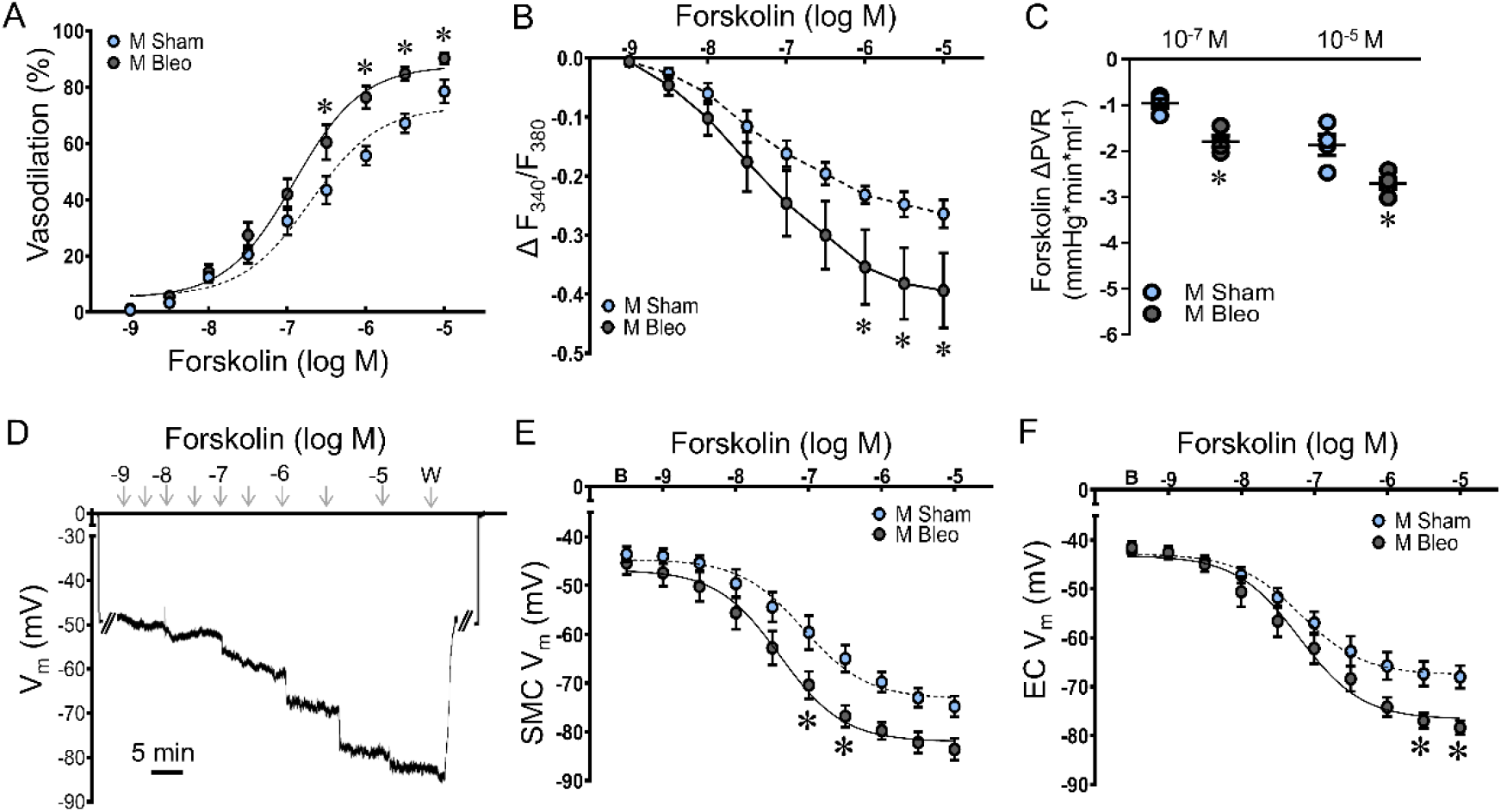
Stimulation of PKA evokes greater vasodilation and hyperpolarization in PAs from PF mice. A) Vasodilation and B) Ca^2+^ responses to forskolin in isolated pressurized PAs from sham and bleomycin treated males. C) Changes in PVR to forskolin (10^-7^-10^-5^ M) in isolated lungs preconstricted with U46619. D) Representative membrane potential (V_m_) recording illustrating hyperpolarization to forskolin in a SMC from a male PA from a bleomycin-treated mouse. Summary data for V_m_ responses to forskolin in E) SMCs and F) ECs. Values are means ± SEM; n=4-5/group. *P<0.05 bleomycin vs. sham.

### Activation of sensory nerves mediates greater CGRP-dependent vasodilation in PAs from PF mice

In order to evaluate if PF alters perivascular sensory CGRPergic innervation of the lung, we performed CGRP staining in isolated PA trees and observed no differences between sham and bleomycin treated mice (Figures 8A and B). Furthermore, sensory nerves were located sparingly throughout the lung sections of both sexes and PF did not alter coverage of CGRPergic nerves (Figure S11).

**Figure 8.**
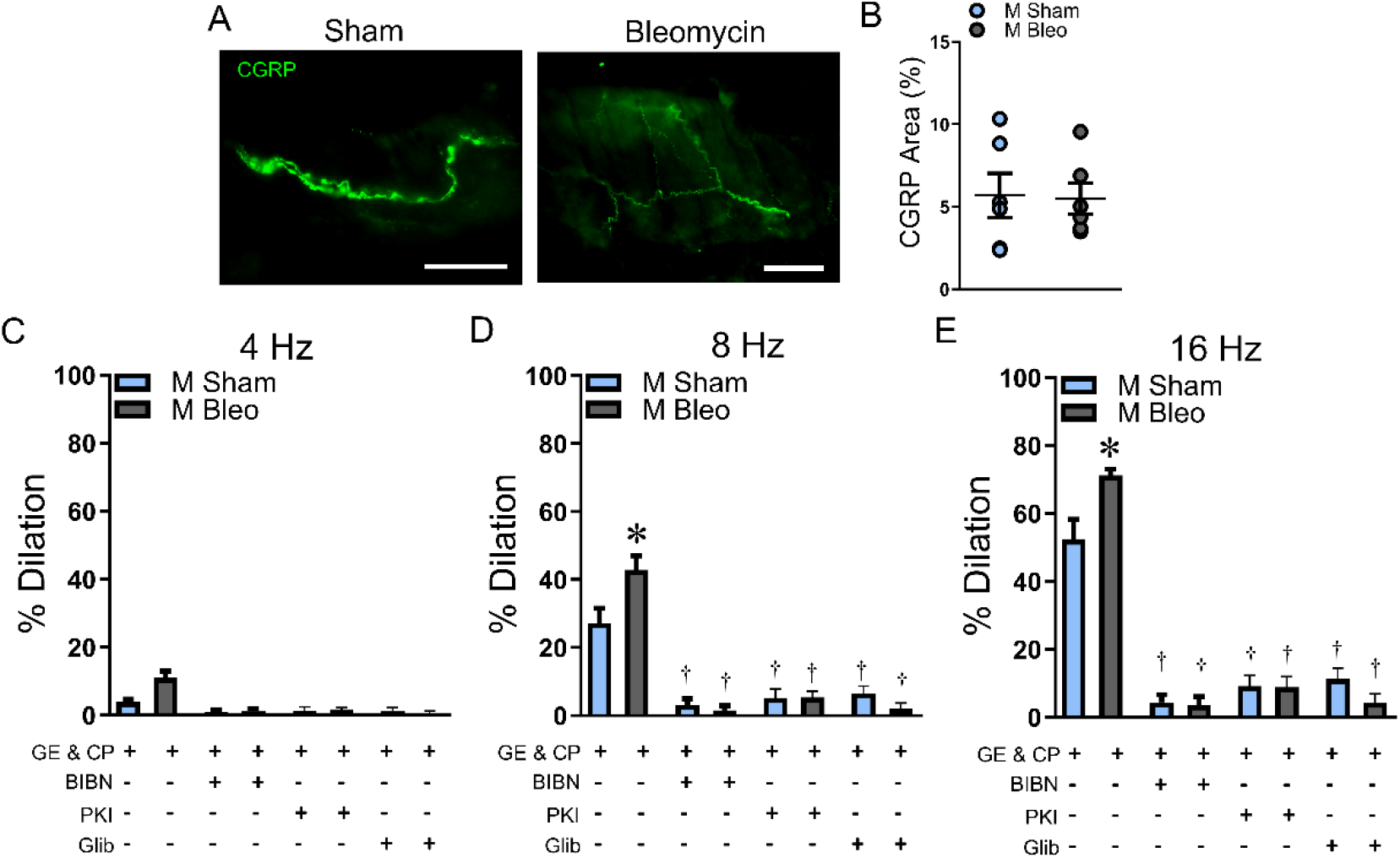
PF augments sensory nerve-mediated dilation. A) Representative CGRP staining on PAs from sham and bleomycin mice; scale bars = 50 µm. B) Summary data for sensory nerve coverage located on PAs. Vasodilation evoked by electric field stimulation (EFS) at C) 4 Hz, D) 8 Hz and E) 16 Hz in PAs from sham and bleomycin mice in the presence or absence of BIBN 4096 (10^-6^ M), PKI (10^-5^ M), and glibenclamide (10^-6^ M). All vessels were treated with the sympathetic blocker guanethidine (10^-5^ M) and the substance P antagonist CP 99994 (10^-6^ M). Values are means ± SEM; n=5-6/group. ^*^P < 0.05 bleomycin vs. sham. ^†^P < 0.05 vs. respective vehicle.

To determine how local activation of sensory nerves contributes to vasodilation, vasodilator responses in isolated, pressurized PAs preconstricted with UTP were evaluated during electrical field stimulation (4-16 Hz)^24^. All vessels were treated with guanethidine and CP 99994 to prevent potential complicating influences of sympathetic nerves and substance P signaling respectively. Electrical stimulation led to frequency-dependent increases in vasodilation in PAs from both sham and bleomycin mice (Figure 8C-E). Vasodilation to electric field stimulation was significantly greater in bleomycin vs. sham vessels and could be prevented by CGRP inhibition, confirming that the actions of perivascular nerve stimulation are CGRP-dependent. Furthermore, blocking K_ATP_ channels or PKA could prevent vasodilation in response to perivascular nerve stimulation consistent with our observations in isolated vessels.

## DISCUSSION

Increasing evidence demonstrates that signaling in PF differs from other mechanisms of hypoxia-induced PH. Given the correlation between the severity of PH during PF and patient outcome^6^, understanding how to prevent or delay the onset of PH represents an important avenue for patient therapy. Whereas, CGRP-induced vasodilator function is compromised in hypoxia-induced PH^15^, these new findings indicate *enhanced* CGRP-induced vasodilator function in PF induced by bleomycin (3 wk). The present data demonstrate that: 1) bleomycin exposure leads to greater PH in males vs. females despite a similar magnitude of fibrosis; 2) CGRP-dependent vasodilation and hyperpolarization was significantly increased in bleomycin treated male, but not female mice; 3) greater vasodilator sensitivity to CGRP is mediated through increased PKA-dependent activation of K_ATP_ channels; and 4) despite similar sensory innervation in PF lungs, perivascular nerve stimulation evokes greater vasodilation in vessels from PF mice. While other studies have identified key differences in vascular structure in PF compared to hypoxia-induced PH^7, 25^, the present data are among the first to evaluate differences in regulation of vascular tone.

### Female Sex Attenuates the Development of PH Induced by PF

Current reports vary as to whether males or females exhibit more severe fibrosis and lung impairments due to bleomycin exposure. These differences may relate to the specific species utilized, age, the time-point studied following bleomycin exposure, or the phase of estrus cycle^26-29^. For the present studies, we observed similar degrees of fibrosis in male and female mice 3 wks following exposure to bleomycin (Figure 1). Female patients with PF have greater survival than males^30^, and this has been suggested to be mediated by greater resilience to hypoxia-induced PH in females. Consistently, we observed reduced indices of PH in mice treated with bleomycin (Figure 1 and Table S1). Differences in location of collagen deposition may relate to disease severity, with males having greater proportion of collagen adjacent to small and intermediate sized airways^26^. Additionally, inflammation can contribute to the development of PH^31, 32^, however the specific contributions of inflammation associated with fibrosis of the lung in this setting of PH and potential sex differences have not yet been defined. Therefore, despite a similar magnitude of fibrosis, the stimuli contributing to PH could be greater in males. For the present studies, we focused on early PF (3 wk bleomycin) to minimize the contribution of hypoxia to focus on differences in signaling between PH initiated by hypoxia and PF.

### PF Augments Vasodilation to CGRP

The therapeutic role for the CGRP pathway is supported by findings that CGRP diminishes the severity of PH^33, 34^ with an inverse correlation between plasma [CGRP] and PA systolic pressure in human patients^35^. While delivery of prepro-CGRP reverses pulmonary vascular remodeling and reduces PA pressure^36, 37^ in chronic hypoxia-induced PH, reduced CGRP availability increases vasomotor tone^15^ and induces vascular remodeling^38, 39^. In contrast, in the setting of PF, CGRP reduces PF^16, 17^. Therefore, we sought to test whether PF would augment vasodilator sensitivity to CGRP. Our findings indicate greater vasodilator sensitivity to CGRP in both isolated lungs and pressurized PAs from males that develop more severe PH (Figure 2) which persisted at elevated intravascular pressures associated with PH. Greater vasodilation in vessels from male PF mice was associated with a greater fall in vessel wall [Ca^2+^]_i_. Given the importance of voltage-gated calcium channels in regulation of vascular tone^40^, we investigated effects of CGRP on V_m_ in PAs. Consistent with differences in vasodilatory function between sexes, we observed that hyperpolarization of SMCs and ECs to CGRP was enhanced in PAs and endothelial tubes from males but not females (Figure 2). Although, females did not exhibit greater vasodilation to CGRP, unlike hypoxia-induced PH, responsiveness was not reduced. Estrogen can reduce CGRP levels in neurons^41^ and CGRP-induced vasodilation is impaired in human middle meningeal arteries from females^42^. Whether such sex differences promote receptor desensitization to CGRP in PAs during PF remains to be fully elucidated.

In the healthy lung, endothelial-derived NO plays a minimal role in vasodilation to CGRP^14^. We sought to evaluate whether endothelial function contributes to enhanced sensitivity to CGRP. Neither endothelial disruption nor inhibition of eNOS prevented the augmented V_m_ hyperpolarization and vasodilation to CGRP in males (Figures S5 and S6), indicating that enhanced sensitivity to CGRP is inherent to the SMC layer of the vessel wall. Nonetheless, although ECs do not experience greater [Ca^2+^]_i_ responses during PF (Figure S7), they do experience a modest increase in hyperpolarization of V_m_ (Figure 2). However, these findings do not preclude a potential role for endothelium-dependent hyperpolarization to provide a modest contribution in this setting. Taken together, these data indicate independent, yet complementary, roles for ECs and SMCs to potentiate vasodilation to CGRP, whether by sensory nerves adjacent to SMCs or when presented to ECs via the circulation.

### Mechanisms of Enhanced Vasodilation to CGRP

Pulmonary vasodilation to CGRP is mediated by PKA-dependent activation of K_ATP_ channels^14^. K_ATP_ channels in the pulmonary vasculature are comprised of four inward rectifying subunits (K_IR_6.1) each associated with a sulfonylurea receptor (SUR2B)^43^. Blocking K_ATP_ signaling pharmacologically with glibenclamide or genetically via K_IR_6.1 knockout nearly abolished hyperpolarization and vasodilation to CGRP in PAs from both sham- and bleomycin-treated male mice eliminating differences between groups (Figures 3 and 4). Furthermore, the RVSP and Fulton index were enhanced in K_IR_6.1 knockout mice administered bleomycin, indicating that this pathway limits the development of PF-induced PH *in vivo*. While K_Ca_ channels do not play a role in pulmonary vasodilation to CGRP under normal conditions^14^, they contribute to CGRP dependent hyperpolarization and vasodilation in systemic arteries^44^. Therefore, we sought to determine whether a role for K_Ca_ channels is unmasked during PF to enhance sensitivity to CGRP. However, neither vasodilation nor V_m_ hyperpolarization were altered by K_Ca_ inhibition (Figures S8 and S9). Next, we utilized pinacidil to evaluate whether direct stimulation of K_ATP_ channels led to greater hyperpolarization and vasodilation in male PF mice. Hyperpolarization of V_m_ with pinacidil was similar between sham and bleomycin treated mice (Figure 5), although the maximal magnitude of hyperpolarization was ∼10 mV greater for pinacidil vs. CGRP in sham vessels suggesting that CGRP does not maximally activate K_ATP_. In agreement with our electrophysiological data, we did not observe differences in vasodilator sensitivity to pinacidil in either pressurized PAs or isolated lungs. Thus, it appears that differences in coupling of CGRP receptors to K_ATP_ activation are what mediate greater vasodilation to CGRP in PAs from males.

PKA can phosphorylate SMC K_ATP_ channels at three residues (S385 for K_IR_6.1 and T633 and S1465 for SUR2B) to facilitate hyperpolarization^45^. We observed that PKA inhibition greatly attenuated CGRP-dependent pulmonary arterial hyperpolarization and vasodilation in vessels and lungs from sham and bleomycin mice (Figure 6). Furthermore, we observed that direct activation of PKA led to greater vasodilation in PF vs. sham mice (Figure 7). This was associated with both enhanced hyperpolarization and a greater fall in vessel wall [Ca^2+^]_i_. While these data clearly indicate an enhanced role for PKA signaling in bleomycin treated lungs from males, whether this results from a change in expression or rather solely through changes in regulation of intracellular signaling domains, remains to be addressed. Furthermore, targeting the cAMP/PKA may provide synergistic effects to limit the severity of fibrotic injury in interstitial lung disease given its ability to limit activation of the NLRP3 inflammasome and pyroptosis in alveolar epithelial cells^46^. Indeed, recent studies have shown that stimulating the PKA pathway with cAMP-containing extracellular vesicles can improve outcomes in rats with hypoxia-induced PH^47^, indicating the therapeutic potential for this pathway in targeting lung disease.

Expression of K_V_ channels decreases in hypoxic pulmonary hypertension^48^ and K_V_ currents can be inhibited by hypoxia or endothelin-1^49, 50^. K_V_ dysfunction is also observed in pulmonary arterial myocytes from patients with primary pulmonary hypertension^51^. The ensuing depolarization elevates SMC [Ca^2+^]_i_ through activation of voltage-gated calcium channels to facilitate sustained vasoconstriction and vascular remodeling^52^. In contrast to chronic hypoxia where SMC V_m_ is depolarized under baseline conditions^53^, we observed no significant effect of PF on resting V_m_. PKA can also phosphorylate and modulate the activity of K_V_ channels promoting hyperpolarization in cerebral vessels^54^. In addition, depolarization of V_m_ leads to enhanced Ca^2+^ sensitization in PH resulting from chronic hypoxia^55^, and PKA can phosphorylate the regulatory subunit (MYPT-1) of myosin light chain phosphatase to prevent Ca^2+^ sensitization^56^. Thus, enhanced PKA signaling in PF may limit enhanced vasoconstriction in PF through mechanisms beyond K_ATP_ function.

### Physiological Stimulation of Sensory Nerves Evokes Greater Vasodilation in PF Arteries

Two pools of CGRP can activate CGPR receptors on pulmonary vessels. The first is from perivascular nerves which run along the adventitia and are in proximity to SMCs^14^. We observed no difference in either total lung or perivascular sensory innervation (Figures 8 and S11). While these findings suggest that CGRP produced within the lung is not altered by PF, they do not preclude potentially greater release of CGRP from nerve terminals present next to the circulation as can occur in the central nervous system ^57^. Circulating CGRP represents the second pool of CGRP that can act on ECs^35^. Presently, conflicting reports exist as to whether CGRP levels are reduced or increased in patients with interstitial lung disease^58, 59^. Regardless of source, receptors for CGRP are composed of CRLR, RAMP1 and receptor component proteins^60^ which are expressed on both pulmonary vascular SMCs and ECs^14^. Despite the reduced sensitivity to CGRP in hypoxia-induced PH, vascular CGRP receptor levels are elevated^61^. Whether there are differences in RAMP1 or CRLR levels in the setting of PF remains to be evaluated, although potential changes in receptor expression would not alter the greater ability for PKA to mediate pulmonary dilation.

While the density of sensory fibers and CGRP receptors play an important role in determining the effects of CGRP, the location of these components in the vessel wall can also play a key role. Due to increased adventitia, large arteries have larger diffusion distances (> 500 nm) than small arteries (∼100 nm), increasing the time to response and reducing the effectiveness of the released chemical concentration^62^. In the setting of lung fibrosis, there may be increased adventitial thickness^63^ that may act to limit the effectiveness of CGRP released *in situ*. However, when perivascular nerves adjacent to isolated PAs are stimulated by an electric field, we observed greater vasodilation (Figure 8) indicating that changes vasodilation persist regardless of alterations in matrix deposition.

While the present investigation focuses on CGRP signaling, sensory nerves also release Substance P. Substance P has been linked to both pulmonary vasoconstriction and vasodilation depending on the species and preparation^64, 65^. While the effects of lung fibrosis on Substance P signaling have not been characterized, Substance P levels are increased in other animal models of PH contributing to development of disease^62^. In addition, endothelial-dependent dilation to CGRP is absent in patients with PH^15^. Signaling crosstalk between various neurotransmitters may further complicate the present findings. In mesenteric arteries from mice with inflammatory bowel disease, Substance P diminishes CGRP-mediated sensory vasodilation^24^. Furthermore, larger species, such as humans often have more sympathetic and parasympathetic nerves in the lung compared to mice^66^. Therefore, future investigations are necessary to fully understand how PF alters the balance of sensory neurotransmitters in the context of PF.

### Conclusion

Studies of group 3 PH have mainly focused on hypoxia-induced PH, failing to account for how differences in signaling which may accompany interstitial lung disease. The current study highlights distinct differences in vascular responses to the sensory neurotransmitter CGRP between early-stage PF and hypoxia-induced PH. Furthermore, we identified key differences in the regulation of vascular tone between males and females. We conclude that CGRP evokes greater pulmonary vasodilation through PKA-dependent activation of K_ATP_ channels in males without altering vasodilator sensitivity in females. Given the presence of PH in PF significantly increases mortality^4^, understanding how to delay the development of PH in PF has the potential to lead to targeted vasodilator therapies that extend survival and improve quality of life in afflicted patients.

## NOVELTY AND SIGNIFICANCE

### What is known?

- Calcitonin gene-related peptide (CGRP) can limit the severity of pulmonary fibrosis (PF), but effects on the vasculature in this disease setting are poorly defined.
- Pulmonary vasodilation to CGRP is mediated by protein kinase A (PKA) and ATP-sensitive potassium channels (K_ATP_).

### What new information does this article contribute?

- Pulmonary hypertension induced by PF is more severe in male vs. female mice.
- Pulmonary arteries from males exhibit more pronounced vasodilation to CGRP, mediated through enhanced activation of K_ATP_ by PKA.
- Although PF does not change perivascular sensory innervation, electric field stimulation evokes greater pulmonary vasodilation in pulmonary arteries from male mice with PF.

### Summary

While studies of group 3 PH have focused on hypoxic mechanisms, vascular dysfunction in interstitial lung diseases has received far less attention. This research provides the first characterization of sensory nerve-mediated changes in the regulation of pulmonary arterial tone during early PF. PH in a bleomycin mouse model of PF is more severe in males vs. females as characterized by greater increases in RVSP and Fulton index. In contrast to hypoxia-induced PH, PF did not reduce vasodilator sensitivity to CGRP and enhanced vasodilation in isolated lungs and pressurized pulmonary arteries. Furthermore, stimulation of sensory nerves evoked a more robust dilation in vessels from male mice with PF. Enhanced vasodilation to CGRP was associated with enhanced PKA-dependent K_ATP_ activation and greater hyperpolarization of smooth muscle cells. These findings highlight the potential for CGRP and PKA pathways to reduce pulmonary vascular resistance and limit the severity of PH associated with PF and suggest that treatment targets may differ for group 3 PH associated with interstitial lung diseases.

## ACKNOWLEDGEMENTS

The author thanks Erin Hoover for technical assistance with bleomycin mice.

## SOURCES OF FUNDING

This research was supported by American Heart Association Career Development Award (CDA931652).

## DISCLOSURES

The authors have no conflicts of interest to disclose.

## NON-STANDARD ABBREVIATIONS AND ACRONYMS

BK: large conductance calcium activated potassium channel
CGRP: calcitonin gene-related peptide
CRLR: calcitonin receptor-like receptor
EC: endothelial cell
eNOS: endothelial nitric oxide synthase
IK: intermittent conductance calcium activated potassium channel
K_ATP_: ATP sensitive potassium channel
K_Ca_: calcium activated potassium channel
K_V_: voltage gated potassium channel
PA: pulmonary artery
PAP: pulmonary arterial pressure
PF: pulmonary fibrosis
PH: pulmonary hypertension
PKA: protein kinase A
PKG: protein kinase G
PVR: pulmonary vascular resistance
RAMP1: receptor activity-modifying protein 1
RV: right ventricle
RVSP: right ventricular systolic pressure
SK: small conductance calcium activated potassium channel
SMC: smooth muscle cell
UTP: uridine triphosphate
V_m_: membrane potential

## Notes

### Competing Interest Statement

The authors have declared no competing interest.

### Summary of Updates

Addition of Supplemental Material including methods, tables, and figures.

