## Supplemental Methods, Tables, and Figures for "Pulmonary Fibrosis Enhances Vasodilation to Calcitonin Gene-Related Peptide"

Charles E. Norton

### DETAILED METHODS

*Animal care and use*

All protocols and experimental procedures were reviewed and approved by the University of Missouri Animal Care and Use Committee (Protocol #40469) and comply with ARRIVE guidelines. Experiments were performed on male (n=268) and female (n=44) C57Bl/6J mice purchased from Jackson Laboratories (4-6 mo old). $K_{ATP}$/ $K_{IR}6.1$ knockout ($K_{ATP}^{-/-}$; mice lacking KCNJ8 gene on C57BL/6 background) mice[1] obtained from Dr. Michael Davis (University of Missouri) were used for complementary experiments. Prior to use, mice were housed on a 12:12 h light dark cycle at ~23°C with fresh water and food available *ad libitum*. Mice were anesthetized with ketamine and xylazine (100 $mg \cdot kg^{-1}$ and 10 $mg \cdot kg^{-1}$ respectively; intraperitoneal injection) to harvest tissue and killed by exsanguination.

*Bleomycin Model of Pulmonary Fibrosis*

Intratracheal delivery of bleomycin is an established and frequently used model of experimental PF[2]. Mice are anesthetized with isoflurane, then placed in a supine position on a surgical board. A rubber band is positioned around the top front teeth and gently retracted to open the mouth. An oropharyngeal tongue pull is used to ensure that the mouse will inhale the agent, as holding the tongue blocks the swallowing reflex. PF is induced via a single 50 µL bolus of bleomycin (containing 0.025 U; Cat. #B5507, Sigma-Aldrich, St. Louis, MO, USA)[3] diluted in normal saline in the back of the throat with a 24G gavage needle. Mice are placed on a heating pad for recovery from anesthesia and studied 3 wk (±2 d) following bleomycin inhalation. The resulting lung fibrosis (Figure 1) with alternating zone of fibrosis and intervening patches of normal lung is consistent with observations in human patients[4]. Sham lung injuries (saline injections) serve as experimental controls and have responses similar to naïve mice[5].

To evaluate the model of PF, we evaluated collagen deposition and cellularity in fixed lung sections. Lungs were exposed by a median sternotomy and heparin (100 U in 0.1 mL) was injected directly into the right ventricle. The pulmonary artery was cannulated through an incision in the right ventricle and perfused with 20 mL physiological salt solution [PSS, pH 7.4; containing (in mM): 140 NaCl (Thermo Fisher Scientific; Waltham, MA, USA), 5 KCl (Fisher), 1 $MgCl_2$ (Sigma), 10 HEPES (Sigma), 2 $CaCl_2$ (Fisher) and 10 glucose (Fisher)] containing papaverine ($10^{-4}$ M; )[6] to maximally dilate vessels in the lung. An incision was made in the left ventricle to allow efflux of perfusate. After wash with PSS, lungs were perfused in fixative [0.1 M phosphate-buffered saline (PBS) with 4% paraformaldehyde and $10^{-4}$ M papaverine (Cat. #P3510, Sigma). Lungs were additionally inflated with fixative via the trachea and then immersed in fixative. Fixed lungs were submitted to IDEXX BioAnalytics Lab (Columbia, MO, USA) for paraffin embedding, sectioning, and labeling with Masson's trichrome stain. Lung sections were examined with DS-Qi2 camera on an Eclipse 800 microscope (both from Nikon; Tokyo, Japan) at 10× (N.A. 0.3) and 40× (N.A. 0.8) magnification. Collagen content was quantified as the % area stained by aniline blue in Masson's trichrome sections[7]. Cellularity (# cells/mm) was determined by counting individual nuclei in 100 µm × 100 µm regions of lung tissue[8].

*Indices of Pulmonary Hypertension*

Peak right ventricular systolic pressure (RVSP) was measured as an index of pulmonary arterial pressure in anesthetized mice[6]. Following an upper transverse laparotomy, a 25G needle was inserted through the diaphragm into the right ventricle. Data were recorded with an APT300 pressure transducer (Harvard Apparatus, Holliston, MA, USA) using Ponemah Pulmodyn software (version 5.60). Fulton's index is expressed as right ventricle to left ventricle + septum weight after removing atria and surrounding connective tissue from the heart. Hematocrit (% red blood cells) was measured in blood samples collected in glass microcapillary tubes after direct cardiac puncture at the time of tissue collection as an index of polycythemia[6].

*Preparation of isolated pulmonary arteries and endothelial tubes*

Individual PAs [~2 mm length, 100-150 µm inner diameter (ID)] were dissected from surrounding lung tissue while viewing through a stereomicroscope in PSS. PAs were cannulated onto heat-polished micropipettes (outer diameter, ~100 µm), secured with silk suture, and placed in a tissue chamber (RC-27N; Warner Instrument; Hamden, CT, USA) and superfused at 3 mL $min^{-1}$. Vessels were pressurized to 16 $cmH_2O$ (~ 12 mmHg) and maintained at 37°C[5]. Once pressurized, any vessels with apparent leaks were discarded.

*Endothelial disruption*. To selectively test SMC responses independent of endothelial influences, the endothelium was disrupted by rubbing a small tungsten wire (50 µm diameter; Goodfellow; Huntington, UK) through the vessel lumen three times prior to cannulating the second end of the vessel. Once the vessel was cannulated and pressurized, selective damage of ECs was verified by constriction to 5 µm uridine triphosphate (UTP; Cat. #U6625, Sigma) with loss of dilation to 10 µM acetylcholine (ACh; Cat. #A6625, Sigma)[5].

*Endothelial tube isolation*. Intact endothelial tubes (devoid of SMCs) were obtained by enzymatically digesting isolated PAs with 62 µg/ml papain (Cat. #P4762, Sigma), 1.0 mg/ml dithioerythritol (Cat. #D8255, Sigma), 1.5 mg/ml collagenase (Cat #C8051, Sigma), and 30 µg/ml elastase (Cat. #LS002290, Worthington Biochemical, Lakewood, NJ) in PSS for 25 min at 32°C[5]. After incubation, the enzyme solution was replaced with normal PSS and arterial segments were transferred to the tissue chamber. During visual observation at 200× magnification, segments were gently triturated to remove SMCs. Trituration pipettes were pulled from borosilicate glass capillary tubes (Cat. #1B100-4, World Precision Instruments; Sarasota, Fl), heat-polished to a tip diameter of ~120 µm, and connected to a Nanoliter injector (World Precision Instruments) for controlled aspiration and ejection of the PA segment. After removal of SMCs, endothelial tubes were secured to the tissue chamber with glass pinning pipettes (heat-blunted tips, ~80 µm). Endothelial tubes were not pressurized and studied at 32°C to maintain stability for experiments[5].

*Vasomotor Responses and $Ca^{2+}$ photometry*

Intact or endothelium-disrupted pressurized PAs were placed on the stage of a Nikon Eclipse TS100 inverted microscope for simultaneous measurement of PA diameter and $[Ca^{2+}]_i$. Using bright-field illumination, images of a PAs were acquired with a 20× objective (N.A. 0.45; Nikon Fluor 20) and a charge-coupled device camera (MyoCam-S, IonOptix, Milford, MA) and diameter was recorded with IonWizard software (version 6.3, IonOptix). For measurement of $[Ca^{2+}]_i$, PAs were incubated in Fura 2-AM dye [dissolved in DMSO and diluted to 1 µM in PSS (final [DMSO] = 0.5%); Cat. #F14158, Fisher] for 40 min without flow. Superfusion with PSS was then resumed for 20 min to wash out excess dye. Fura 2 fluorescence was used to evaluate $[Ca^{2+}]_i$ by alternatively exciting the preparation at 340 and 380 nm while recording emissions at 510 nm through a 20× Fluor 20 Nikon objective (N.A. = 0.45) using IonWizard 6.3 software. Under these conditions, Fura 2 fluorescence primarily reflects SMC $[Ca^{2+}]_i$[5]. During focus on the widest section of the vessel, ID and $[Ca^{2+}]_i$ was recorded during the $5^{th}$ min at each concentration of agonist using a personal computer. After subtracting background fluorescence (F), $[Ca^{2+}]_i$ values are expressed as $F_{340}/F_{380}$ ratios. Concentration response curves to UTP ($10^{-9}$-$10^{-3}$ M) provided an $EC_{50}$ ($5\times10^{-6}$ M) to induce a stable pressor response in vessels and examine dilation to CGRP. Concentration response curves to CGRP ($10^{-10}$-$10^{-6}$ M; Cat. #AS-20681, AnaSpec; Freemont, CA) were performed in half-log molar steps, each for 5 min, to evaluate its effect on arterial diameter and $[Ca^{2+}]_i$ in PAs from sham and PF mice. The contribution of nitric oxide (NO) was evaluated with the NO synthase inhibitor L-NAME ($10^{-4}$ M; Cat. #N5751, Sigma)[5]. These experiments were repeated in the presence of glibenclamide (1 µm; Cat. #G2539, Sigma) to inhibit $K_{ATP}$ channels[5] in addition to vessels from $K_{ATP}^{-/-}$ mice. The role of $K_{Ca}$ channels was evaluated with charybdotoxin to inhibit large and intermediate conductance channels ($1\times10^{-7}$ M; Cat. #28244, AnaSpec) + apamin to inhibit small conductance channels ($3\times10^{-7}$ M; Cat. #60772, AnaSpec)[9]. Protein kinase inhibitor [$1\times10^{-5}$ M; PKI-(14-22) amide, Cat. #2546; Tocris Bioscience; Bristol, UK][10] was used to evaluate the contribution of PKA; all preparations were equilibrated for 20 min with respective inhibitors prior to evaluation of vasomotor or electrophysiological responses. Concentration response to the $K_{ATP}$ agonist pinacidil ($10^{-9}$-$10^{-5}$ M; Cat. #P154, Sigma)[5] and the PKA agonist forskolin ($10^{-9}$-$10^{-5}$ M; Cat. #F6886, Sigma)[11] were performed in the same manner as for CGRP.

*Endothelial tubes*. For evaluation of $[Ca^{2+}]_i$ responses in the endothelium, freshly isolated endothelial tubes were incubated for 30 min with Fura 2 and rinsed with PSS for 15 min to remove excess dye. Endothelial $[Ca^{2+}]_i$ responses were measured in response to CGRP ($10^{-10}$-$10^{-6}$ M) in endothelial tubes from sham and PF mice.

*Perivascular nerve stimulation*. The effects of direct nerve activation on vasodilation to CGRP were evaluated by electrical field stimulation of isolated, pressurized PCAs. An electrical field was generated by placing the vessel between two platinum electrodes connected to a stimulation isolation unit (SIU5, Grass) and a square wave stimulator (S88, Grass). To measure sensory vasodilation in PAs preconstricted with UTP, electrical pulses (70 V, 2 ms) were delivered at 4, 8, and 16 Hz until a stable response was achieved (~30 s). To selectively evaluate CGRP-dependent dilation PAs were treated with the sympathetic blocker guanethidine ($10^{-5}$ M; Cat. #645-43-2, BOC Biosciences, Frankfurt, Germany) and the neurokinin 1 (substance P) receptor antagonist CP99994 ($10^{-6}$ M; Cat. #3419, Tocris)[12]. To verify responses were mediated by CGRP, vessels were treated with the CGRP receptor antagonist BIBN 4096 ($10^{-6}$ M; Cat. #4561, Tocris)[12]. Experiments were repeated in the presence of glibenclamide and PKI to evaluate the contributions of $K_{ATP}$ channels and PKA respectively to dilation evoked by electric field stimulation.

*Isolated Lung Protocols*

Mice from each group were anesthetized with ketamine/xylaxine, the trachea was cannulated to provide constant positive-pressure ventilation (80 breaths/min, tidal volume 150 mL) with a Harvard rodent ventilator (Minivent 845, Harvard Apparatus)[13] containing a humidified gas mixture composed of 21% $O_2$, 5% $CO_2$, 74% $N_2$. Positive end-expiratory pressure was set at 3 $cmH_2O$ and tracheal pressure ($P_{Trach}$) were recorded with a pressure transducer (DLP2.5 Harvard Apparatus). Mice were placed in the chamber of the isolated perfused lung system (IL-1 839, Harvard Apparatus), After the anterior chest plate was excised and discarded, the pericardium, thymus, and fat tissue were carefully removed. To prevent coagulation, heparin

(20 units) was injected directly into the RV, and the pulmonary artery was cannulated and secured with a 6-0 silk suture. The preparation was immediately perfused with bicarbonate buffered PSS (replacing HEPES with 19 mM $NaHCO_3$, Sigma) containing 4% (wt/vol) albumin (Cat. #BP9703, Sigma) and meclofenamate (30 μM; Sigma) using a peristaltic pump (P70, Harvard Apparatus). Next, a stainless-steel cannula was inserted into the left atrium and fixed in place with a 6-0 silk suture. After rinsing out blood from the lung, perfusate volume was maintained at 50 mL and the flow rate was maintained at 2 mL/min[13]. Pulmonary arterial (PAP) and left ventricular pressure were measured using APT300 pressure sensors and Ponemah Pulmodyn software. To maintain humidity and temperature (37°C) preparations were enclosed with the water jacketed lid for the IL-1 system.

After a 20-min period to stabilize and measure baseline pressure, the thromboxane analog 9,11-dideoxy-11α, 9α-epoxymethanoprostaglandin $F_{2\alpha}$ (U-46619; Cat. #16450 Cayman Chemical) was added to the perfusate reservoir until a stable arterial pressor response of ~10 mmHg was achieved. U-46619 provides consistent and stable pressor responses in this preparation[14, 15], allowing assessment of subsequent vasodilatory responses. To examine the effects of PF on CGRP-dependent pulmonary vasodilation, a cumulative dose-response relationship to CGRP ($10^{-8}$ and $10^{-6}$ M) was assessed in lungs from bleomycin and sham treated mice. To determine whether PF augments $K_{ATP}$- or PKA-mediated vasodilation, cumulative dose-response relationships to pinacidil ($10^{-7}$ and $10^{-5}$ M) and forskolin ($10^{-7}$ and $10^{-5}$ M) were assessed in U-46619 preconstricted lungs from both groups. A stable vasodilatory response to the first dose of each vasodilator was allowed to develop (~10 min) before administration of the second dose.

*Intracellular Recording of Membrane Potential $Ca^{2+}$ photometry*

The $V_m$ of SMCs of intact and endothelium-disrupted PAs and ECs of endothelial tubes was recorded with an AxoClamp 2B amplifier (Molecular Devices; Sunnyvale, CA) using sharp microelectrodes pulled (P-97, Sutter Instruments; Novato, CA, USA) from glass capillary tubes (Cat. #GC100F-10, Warner) backfilled with 2 M KCl (~150 MΩ tip resistance). A Ag/AgCl pellet in the superfusate was used as a reference electrode. The output amplifier was connected to a data acquisition system (Digidata 1322A; Molecular Devices) and an ABM-3 audible monitor (World Precision Instruments). Successful impalements of SMCs and ECs were indicated by: 1) sharp negative deflection of $V_m$, 2) stable $V_m$ after impalement (>1 min), 3) depolarization to 100 mM KCl (SMCs) or hyperpolarization to 10 μM ACh (ECs), 4) recovery to resting $V_m$ after washout of pharmacological agents, and 5) prompt return to ~0 mV upon withdrawal of electrode[5]. Data were acquired at 1 kHz with AxoScope 10.1 software (Molecular Devices). After cell impalement, baseline $V_m$ was recorded for 10 min to ensure stable electrode placement. Increasing concentrations of CGRP ($10^{-10}$-$10^{-6}$ M) were applied to PAs and endothelial tubes for 5 min intervals. Data were averaged for 30 s during stable $V_m$ at each [CGRP] for analysis. Experiments were repeated in PAs from $K_{ATP}^{-/-}$ mice, and vessels pretreated with glibenclamide or PKI. Concentration response to pinacidil ($10^{-9}$-$10^{-5}$ M) was used as a measure of $K_{ATP}$ function and forskolin ($10^{-9}$-$10^{-5}$ M) was used as a measure of PKA function in sham and PF mice.

*Immunofluorescence for CGRP-Positive Nerves in Lung Tissue*

The upper right lobe of lungs from sham and bleomycin mice were placed in optimal cutting temperature compound (OCT, Cat. #23-730-571, Fisher), frozen in isopentane placed in liquid nitrogen, and stored at -80°C. Sagittal sections (thickness, 14 μm) were obtained using a cryostat (Microm HM550, Thermo Scientific) and placed on Superfrost slides (Cat. #1255015, Fisher). Slides were allowed to dry for ~30 min, tissue sections were circled with a PAP pen, then rinsed with PBS (pH=7.4; Cat. #P3810, Sigma) for 5 min to remove excess OCT. Preparations were incubated for 30 min with picrosirius red (Cat. #365548, Sigma) in picric acid, then rinsed with 0.5% acetic acid, and then ethanol (70%)[16]. Lung sections were blocked for 15 min with 10% normal goat serum (Cat. #566380, Sigma) containing 0.5% Triton X-100 (Cat. #T8787, Sigma) in PBS, then incubated with a rabbit polyclonal CGRP primary antibody (1:250 dilution; Cat. #PC205L, Millipore; Billerica, MA) for 1 h. The selectivity of this antibody has been validated with pre-absorption in mouse spinal cord sections[17]. Preparations were then washed twice with PBS, then incubated for 1h with an Alexa Fluor 488 secondary antibody (1:500 dilution; Cat. #A32731TR, Molecular Probes; Carlsbad, CA). Slides were rinsed three times with PBS then cover slipped with Prolong Diamond (Cat. #P36965, Invitrogen; Carlsbad, CA). Slides were placed on the stage of a Leica (Wetzlar, Germany) DMi8 microscope and illuminated with an LED light source. Images were acquired with a HC PL APO 40×/0.95 objective using a K8 monochrome camera and LASX software using consistent settings for all image acquisitions. Mean CGRP fluorescence area was quantified for each image using ImageJ.

*Immunofluorescence for CGRP-Positive Perivascular Sensory Nerves*

Staining for CGRP identified the presence of perivascular sensory nerves[5]. The PA tree consisting of ≥80 μm diameter arteries dissected from the left lobe of the lung and pinned to transparent silicone elastomer (Sylgard 184; Dow Corning, Midland, MI, USA) in 12-well plate. Preparations were fixed in 4% paraformaldehyde for 60 min then washed with phosphate-buffered saline (PBS). Arterial trees were incubated with 10% goat serum and 0.2% Triton X-100 in PBS and then treated for 60 min at room temperature with a CGRP primary antibody (1:250)[17]. Following rinse in PBS, arterial trees were then incubated for 60 min with an Alexa Fluor 488 secondary antibody (1:500 dilution) and washed again in PBS. Arterial trees were mounted on slides with ProLong Diamond under a coverslip then imaged with a 40× objective on a DMi8 microscope (Leica). Sensory nerve density was determined by quantification of fluorescent area of CGRP staining on maximal-intensity z-stack projections of PA segments[18]. Background fluorescence from a region of vessel not containing a nerve fiber was subtracted, the remaining pixels were counted and expressed as a percentage of the total number of pixels in the vessel area.

*Data Analysis and Statistics*

Vasomotor responses are presented as percent constriction or dilation calculated as follows: Vasoconstriction (%) $[(ID_{base} - ID_{resp})/ID_{base}] \times 100$, and Vasodilation (%) = $[(ID_{resp} - ID_{UTP})(ID_{base} - ID_{UTP})] \times 100$, where $ID_{base}$ is baseline ID under control conditions before constriction with UTP, $ID_{resp}$ is response ID during a given agonist concentration, and $ID_{UTP}$ is ID of vessel preconstricted with UTP ($EC_{50}$) before dilator application. For isolated lung experiments, total pulmonary vascular resistance (PVR) was calculated as the difference between pulmonary arterial and pulmonary venous pressures ($P_a$ and $P_v$ respectively) divided by flow. Electrophysiological analyses included resting $V_m$ (mV) under control conditions; peak $V_m$ (mV), i.e., the maximal response to pharmacological intervention; and change in $V_m$ ($\Delta V_m$), i.e., peak-response $V_m$ – preceding baseline $V_m$. Student's t-tests or ANOVA (Prism 11, GraphPad Software; La Jolla, CA) were used to analyze data as appropriate. When significant main effects were detected by ANOVA, Bonferroni's post hoc analysis was performed. $P \leq 0.05$ was considered statistically significant.

Summary data are presented as means ±SE; n refers to the number of vessels (each from a different mouse) in each experimental group.

**SUPPLEMENTAL DATA**

**Supplemental Tables**

Table 1. *Body weight, ventricular weight ratios, and hematocrit*

| Group | BW (g) | RV/BW (mg/g) | LV+S/BW (mg/g) | Hct (%) | n |
|---|---|---|---|---|---|
| Male Sham | 33±2 | 0.72±0.04 | 2.5±0.2 | 47±3 | 10 |
| Female Sham | 29±1[#] | 0.75±0.04 | 2.6±.04 | 42±2[#] | 10 |
| Male Bleomycin | 30±1* | 1.32±0.11* | 2.6±0.3 | 58±3* | 10 |
| Female Bleomycin | 28±1 | 1.18±0.08* | 2.7±0.4 | 49±3*[#] | 10 |

Values are means ± SEM. n = number of mice. BW, body weight; RV, right ventricle weight; LV+S, left ventricle plus septum weight; Hct, hematocrit. *$P<0.05$ bleomycin vs. sham. [#]$P<0.05$ female vs. male.

Table 2. *Baseline values for isolated vessel experiments*

| Group | Baseline Diameter (μm) | Baseline $Ca^{2+}$ ($F_{340}/F_{380}$) | $Ca^{2+}$ Free Diameter (μm) | n |
|---|---|---|---|---|
| Male Sham | 129±4 | 0.65±0.04 | 132±4 | 10 |
| Female Sham | 124±6 | 0.59±0.04 | 128±6 | 10 |
| Male Bleomycin | 125±5 | 0.59±0.02 | 133±5 | 10 |
| Female Bleomycin | 126±6 | 0.67±0.05 | 134±6 | 10 |

Values are means ± SEM. n = number of vessels/group. No significant differences.

Table 3. *Isolated Lung Parameters*

| Group | $P_{Trach}$ | U-46619 (nM) | ΔPVR (U-46619) | n |
|---|---|---|---|---|
| Male Sham | 9.3±3.3 | 394±31 | 3.75±0.48 | 4 |
| Female Sham | 9.6±2.2 | 387±48 | 4.03±0.14 | 4 |
| Male Bleomycin | 18.3±4.4* | 325±29 | 4.15±0.45 | 4 |
| Female Bleomycin | 16.5±1.9* | 338±25 | 4.26±0.31 | 4 |

Values are means ± SEM. n = number of mice. $P_{Trach}$, tracheal pressure; ΔPVR, change in pulmonary vascular resistance (mmHg*min*$ml^{-1}$) in response to U-46619. *$P<0.05$ bleomycin vs. sham.

Table 4. *$V_m$ of ECs and SMCs in PAs from male and female mice at rest and in response to CGRP.*

| | $V_m$, mV | | | |
|---|---|---|---|---|
| SMCs | Rest | Max | Δ | $EC_{50}$, nM |
| Male Sham | -44±3 | -65±1 | -21±3 | 15 |
| Female Sham | -43±2 | -62±2 | -20±5 | 15 |
| Male Bleomycin | -43±3 | -66±3 | -24±4 | 2* |
| Female Bleomycin | -43±3 | -65±2 | -21±6 | 16 |
| ECs | | | | |
| Male Sham | -44±3 | -67±4 | -23±4 | 8 |
| Female Sham | -45±4 | -67±3 | -21±5 | 6 |
| Male Bleomycin | -45±4 | -71±4 | -26±2 | 3* |
| Female Bleomycin | -46±3 | -67±4 | -21±6 | 8 |

Values are means ± SEM; n=4-5 vessels/group. $V_m$, membrane potential; Rest, resting $V_m$; Max, $V_m$ at maximum response during exposure to CGRP ($10^{-10}$-$10^{-6}$ M); Δ, Max – Rest; ECs, endothelial tubes; SMCs, smooth muscle cells. *$P<0.05$ bleomycin vs. sham in males.

Table 5. *Changes in $V_m$ with KCl and ACh confirm electrode placement in SMCs and ECs.*

| | $V_m$, mV | | | |
|---|---|---|---|---|
| SMCs (KCl) | Rest | Max | Δ | n |
| Sham Vehicle | -44±2 | -6±6 | 39±2 | 4 |
| Bleomycin Vehicle | -43±2 | -6±6 | -37±3 | 4 |
| Sham Glibenclamide | -46±1 | -9±6 | -38±4 | 4 |
| Bleomycin Glibenclamide | -44±2 | -7±2 | -38±2 | 4 |
| ECs (ACh) | | | | |
| Sham Vehicle | -43±2 | -81±3 | -38±2 | 4 |
| Bleomycin Vehicle | -43±1 | -78±4 | -35±3 | 4 |
| Sham Glibenclamide | -45±2 | -78±3 | -33±2 | 4 |
| Bleomycin Glibenclamide | -45±3 | -80±3 | -36±5 | 4 |

Values are means ± SEM. n = number of vessels/group. $V_m$, membrane potential; Rest, resting $V_m$, after washout at the end of experiment, but prior to application of KCl or acetylcholine (ACh); Max, $V_m$ at maximum response to KCl ($10^{-1}$ M) or ACh ($10^{-5}$ M); Δ, Max – Rest; ECs, endothelial tubes; SMCs, smooth muscle cells. No significant differences.

**Supplemental Figures**

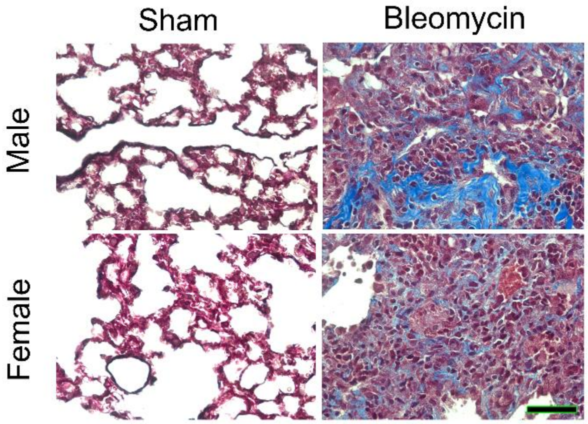


**Supplemental Figure 1. Bleomycin Induces PF**. Representative Masson's Trichrome staining in Sham and Bleomycin treated lungs from male and female mice at 40× magnification (scale bar = 50 μm).

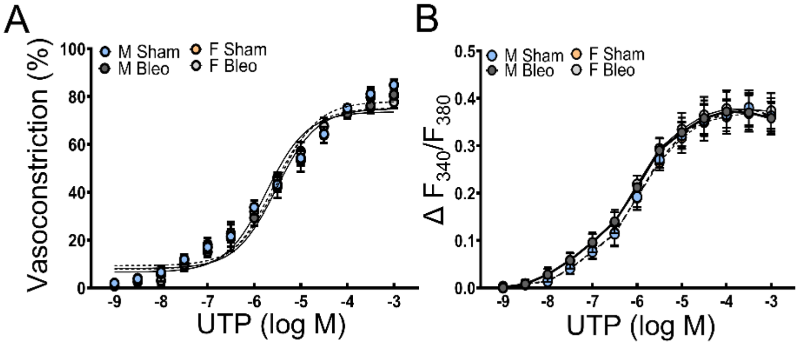


**Supplemental Figure 2. UTP evokes consistent constriction and $Ca^{2+}$ responses in male and female bleomycin and sham mice**. Concentration response curves demonstrating vasoconstriction (A) and corresponding $Ca^{2+}$ responses (B) to UTP ($10^{-9}$-$10^{-3}$ M) in isolated PAs from male (M) and female (F) mice following bleomycin (Bleo) or sham treatment. Values are means ± SEM; n=4 vessels/group. No significant differences were observed.

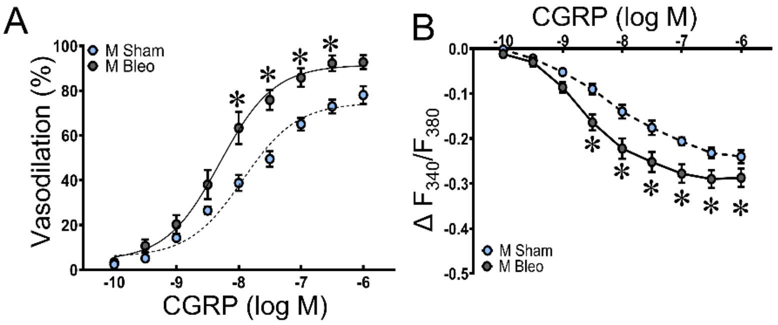


**Supplemental Figure 3. Enhanced vasodilator sensitivity to CGRP persists at elevated pulmonary arterial pressures**. A) Vasodilation and B) corresponding $Ca^{2+}$ responses to CGRP ($10^{-10}$-$10^{-6}$ M) in male PAs pressurized to 40 cm$H_2O$ preconstricted with UTP ($EC_{50}$). Values are means ± SEM; n=5 vessels/group. *P<0.05 bleomycin vs. sham.

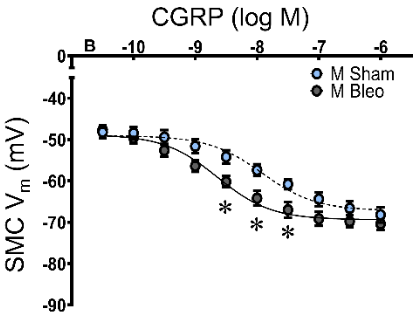


**Supplemental Figure 4. Enhanced hyperpolarization to CGRP persists at elevated pulmonary arterial pressures**. Summary data for $V_m$ of SMCs from intact PAs pressurized to 40 $cmH_2O$ in response to CGRP ($10^{-10}$-$10^{-6}$ M) in PAs from male mice. Values are means ± SEM; n=5 vessels/group. *P<0.05 bleomycin vs. sham.

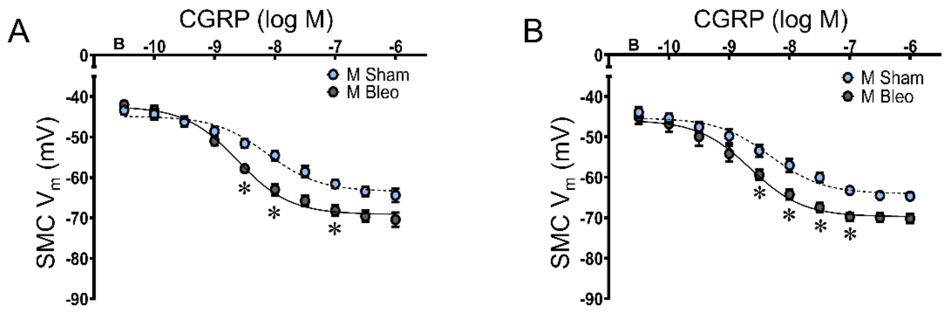


**Supplemental Figure 5. Endothelial nitric oxide signaling does not mediate greater hyperpolarization to CGRP in PF mice**. Summary data for $V_m$ of SMCs from A) endothelium-disrupted PAs and B) PAs treated with L-NAME ($10^{-4}$ M) in response to CGRP ($10^{-10}$-$10^{-6}$ M). Values are means ± SEM; n=5 vessels/group. *P<0.05 bleomycin vs. sham.

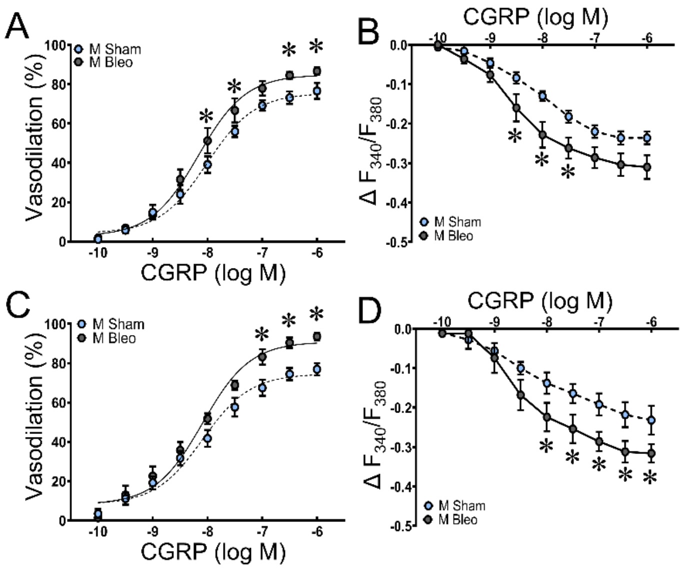


**Supplemental Figure 6. Nitric oxide is not required for enhanced vasodilation to CGRP**. A) Vasodilation and B) corresponding $Ca^{2+}$ responses to CGRP ($10^{-10}$-$10^{-6}$ M) in endothelium disrupted PAs preconstricted with UTP ($EC_{50}$). C) Vasodilation and D) $Ca^{2+}$ responses to CGRP in preconstricted PAs in the presence of L-NAME ($10^{-4}$ M). Values are means ± SEM; n=5 vessels/group. *P<0.05 bleomycin vs. sham.

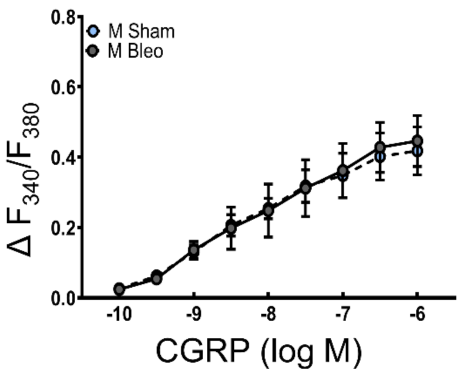


**Supplemental Figure 7. Endothelial [$Ca^{2+}$]$_i$ responses are similar between sham and PF mice**. [$Ca^{2+}$]$_i$ responses to CGRP ($10^{-10}$-$10^{-6}$ M) in endothelial tubes from sham and bleomycin treated mice. Values are means ± SEM; n=5 preparations/group. No significant differences.

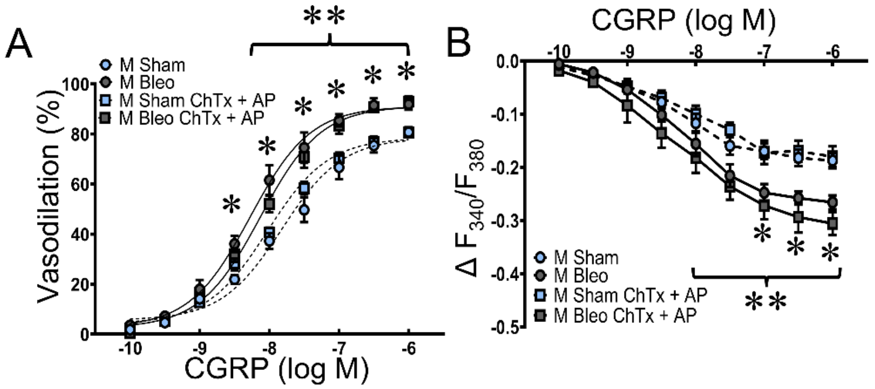


**Supplemental Figure 8. $K_{Ca}$ channels do not mediate greater vasodilation to CGRP in PAs from PF mice**. A) Concentration response curves to CGRP ($10^{-10}$-$10^{-6}$ M) in PAs from sham and bleomycin mice in the absence or presence of charybdotoxin (ChTx; $1\times10^{-7}$ M) and apamin (AP; $3\times10^{-7}$ M). B) Corresponding changes in $[Ca^{2+}]_i$ during exposure to CGRP. Values are means ± SEM; n=5 vessels/group. *P<0.05 bleomycin vs. sham in vehicle. **P<0.05 bleomycin vs. sham in ChTx +AP.

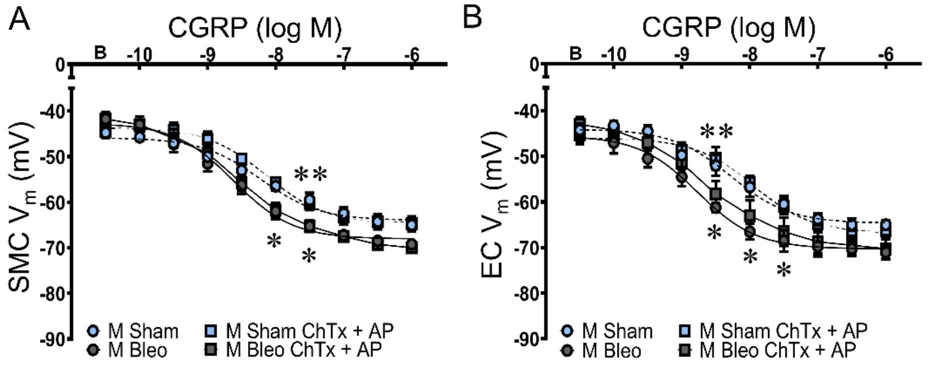


**Supplemental Figure 9. $K_{Ca}$ channels do not mediate enhanced hyperpolarization to CGRP in PAs from PF mice**. Summary $V_m$ data during cumulative exposure to CGRP ($10^{-10}$-$10^{-6}$ M) for A) SMCs of intact PAs and B) ECs from endothelial tubes in the absence and presence of charybdotoxin (ChTx; $1\times10^{-7}$ M) and apamin (AP; $3\times10^{-7}$ M). Values are means ± SEM; n=4 vessels/group. *P<0.05 bleomycin vs. sham in vehicle. **P<0.05 bleomycin vs. sham in ChTx + AP.

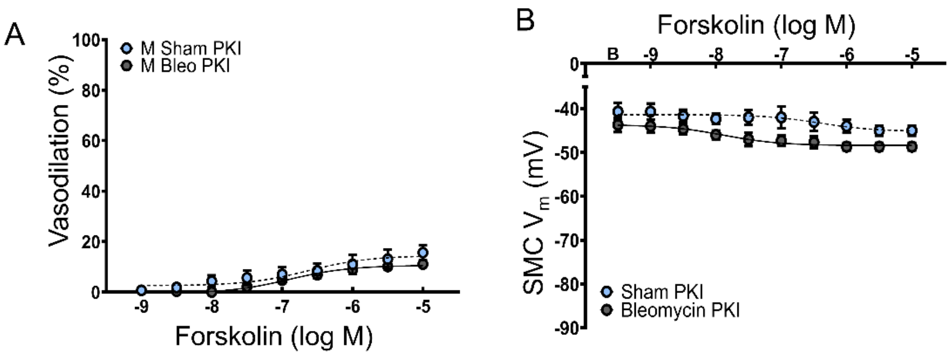


**Supplemental Figure 10. PKA inhibition nearly abolishes vasodilation and hyperpolarization to forskolin in PAs.** A) Vasoconstriction and B) summary $V_m$ data for cumulative exposure to forskolin ($10^{-9}$-$10^{-5}$ M) in intact pulmonary arteries in the presence of PKI ($10^{-5}$ M). Values are means ± SEM; n=3 vessels/group. No significant differences.

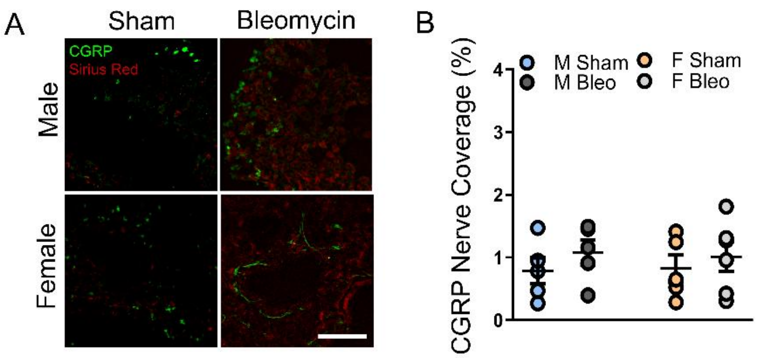


**Supplemental Figure 11. Sensory innervation of the lung is not altered by sex or bleomycin treatment.** A) Representative images and B) summary data for CGRP (indicating sensory nerves) in lung sections counter stained with Sirius red; scale bar = 50 µm. Values are means ± SEM; n=5/group; no statistical differences.
